# Co-option of CENP-A for activity-induced neuronal plasticity

**DOI:** 10.64898/2026.07.16.738894

**Authors:** Ana Stankovic, Daniel Gonzalez-Bohorquez, Izaskun Mallona, Margaux G. Quiniou, Baptiste N. Jaeger, Vladyslav I. Korobeynyk, Linda Brandi, Sandra Günther, Annette E. Rünker, Alexander Garthe, Natalia A. Cruz-Ochoa, Csaba Földy, Scott N. Furlan, Gerd Kempermann, Mark D. Robinson, Sebastian Jessberger

**Affiliations:** Laboratory of Neural Plasticity, Brain Research Institute, University of Zurich, 8057 Zurich, Switzerland; University Research Priority Program (URPP), Adaptive Brain Circuits in Development and Learning (AdaBD), University of Zurich, 8057 Zurich, Switzerland; Department of Molecular Life Sciences, University of Zurich, 8057 Zurich, Switzerland; SIB Swiss Institute of Bioinformatics, University of Zurich, 8057 Zurich, Switzerland; German Center for Neurodegenerative Diseases (DZNE) Dresden and Center for Regenerative Therapies Dresden (CRTD), Technische Universität Dresden, Dresden 01307, Germany; Laboratory of Neural Connectivity, Brain Research Institute, University of Zurich, 8057 Zurich, Switzerland; Ben Towne Center for Childhood Cancer Research, Seattle Children’s Research Institute, the University of Washington, and the Fred Hutchinson Cancer Research Center, Seattle WA 98101, USA

## Abstract

Neuronal activation drives activity-dependent gene expression that underlies experience-associated synaptic modifications, learning and memory. Here we show that the histone variant CENP-A, best known for specifying centromere identity, is dynamically regulated by synaptic activity at both the RNA and protein levels in postmitotic neurons. Neuronal activation increases a non-centromeric nuclear pool of CENP-A while leaving centromeric CENP-A levels unchanged. Downregulation of CENP-A selectively reduces the activity-associated non-centromeric pool, impairs activity-dependent induction of immediate-early genes such as FOS and ARC, and disrupts hippocampus-dependent learning and memory. Furthermore, we find that activity-dependent neuronal responses in human embryonic stem cell-derived forebrain organoids similarly require CENP-A. Our results reveal a mitosis-independent, conserved role of CENP-A for driving plasticity in mammalian neurons.

## INTRODUCTION

Centromeres drive chromosome partitioning during mitosis by nucleating the kinetochore, which acts as an anchor between the mitotic spindle and sister chromatids^1, 2^. Epigenetic specification of a single centromere per chromosome, which is critical for proper chromatid segregation, depends on a unique histone variant called CENP-A. CENP-A epigenetically specifies the location of the centromere, conferring its stable inheritance throughout cell divisions^3–5^. CENP-A is necessary and sufficient for the formation of a functional centromere, partly due to its hyperstability and a dedicated loading machinery allowing for propagation and maintenance of epigenetically defined centromeres^1, 6–11^. Although centromeres and kinetochores are classically viewed as mitosis-specific structures, several kinetochore components are expressed in postmitotic neurons, where they contribute to neurite outgrowth and cytoskeletal organization during neurodevelopment^12–14^. Whether CENP-A itself, or other centromere-associated proteins, play a functional role in mature, postmitotic neurons has remained unexplored. Neurons are excitable cells, capable of generating action potentials, which in turn can drive synaptic changes associated with experience-dependent neuronal modifications. Notably, learning and memory cause changes in synaptic connectivity and neural circuit function, depending on the activity-mediated expression of immediate early genes (IEGs) in activated neurons^15–23^. Given the strict coupling of CENP-A expression to cell division^24–30^, it has been assumed to be largely irrelevant in postmitotic cells. Here, we tested the hypothesis that CENP-A expression and function might be repurposed in neurons as a consequence of neuronal activity. We show that CENP-A is dynamically regulated by synaptic activity in postmitotic neurons, leading to the accumulation of a distinct non-centromeric nuclear pool. This activity-associated CENP-A is required for the full activity-dependent induction of IEGs and hippocampus-dependent learning and memory. Furthermore, neuronal function of CENP-A is conserved between mouse and human neurons. Our findings reveal an unexpected, mitosis-independent role for CENP-A in regulating activity-dependent neuronal plasticity.

## RESULTS

### Neuronal activity drives accumulation of non-centromeric CENP-A

In proliferating cells, CENP-A mRNA expression is tightly regulated across the cell cycle and peaks during G2 phase^24–29^. Surprisingly, in post-mitotic neurons, we observed a rapid upregulation of both Cenp-a mRNA and CENP-A protein following experience-dependent neuronal activation in the mouse hippocampus, a brain region critically involved in specific forms of learning and memory (Figure 1A, Figure 1-figure supplement 1A)^21, 31^. Neuronal activation was induced by exposing mice to a novel environment (NE), which activates a subset of dentate gyrus (DG) neurons, as determined by immunohistochemistry and single-cell RNA sequencing (scRNA-seq) (Figure 1A, 1C)^32^. Cenp-a mRNA dynamics were comparable to those of previously characterized IEGs, such as Fos and Arc (Figure 1A)^17^. Among genes encoding components of the constitutive centromere associated network (CCAN), Cenp-a was uniquely regulated with neuronal activity (Figure 1-figure supplement 1B)^32^. Consistent with this rapid induction upon neuronal activation, the Cenp-a promoter exhibits high chromatin accessibility even in the basal neuronal state, as revealed by previously published ATAC-seq data (Figure 1-figure supplement 1C)^33^. Such constitutive promoter accessibility is a hallmark of a subset of IEGs and confers competence for rapid transcriptional activation^34^. This activity-regulated transcriptional burst is further translated into increased CENP-A protein synthesis rate^17^ (Figure 1-figure supplement 1D) and elevated total protein levels in Pentylenetetrazol (PTZ)-activated DG neurons (Figure 1B, Figure 1-figure supplement 1E)^35^.

**Figure 1.**
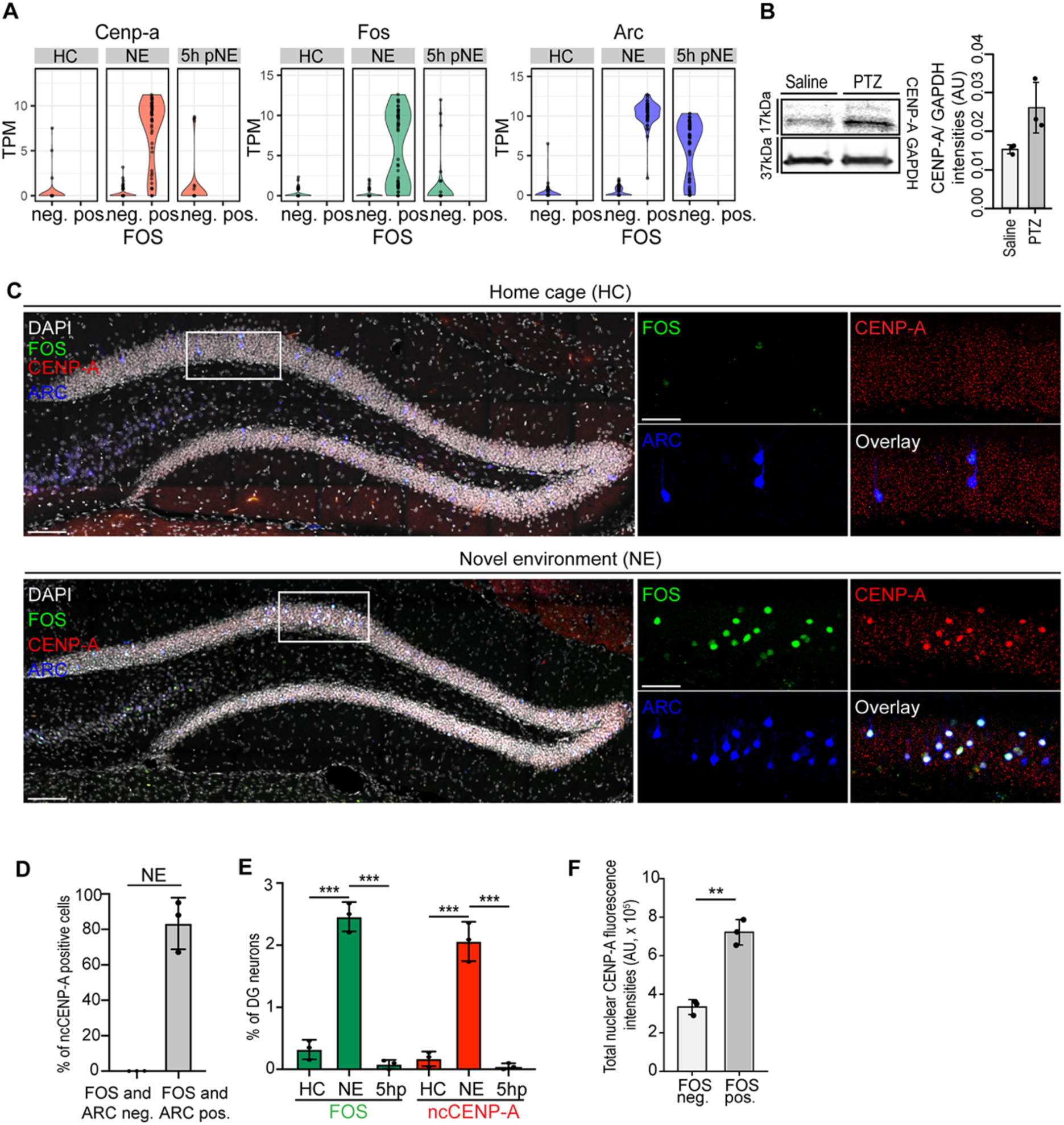
Non-centromeric CENP-A accumulates in activated dentate gyrus neurons. A, Violin plots derived from dataset^32^ showing Fos, Cenp-a and Arc mRNA in homecage (HC), 1 hour (h) after exploring a novel environment (NE), and 5h after NE (5h pNE) using scRNA-seq. nd, not detected. B, Western blot of hippocampal lysates from saline- and PTZ-treated mice, probed for CENP-A and GAPDH, with the CENP-A/GAPDH intensity ratios quantified. CENP-A was elevated in PTZ-treated mice relative to saline controls (mean 1.7-fold; all three PTZ animals exceeded all three saline animals). N=3 mice per group (separate animals per condition). Unpaired two-tailed Welch’s t-test, P=0.10. This tissue-level trend is concordant with the cell-level increase in nuclear CENP-A (F). Data are Mean ± SD. C, Novel environment (NE) exploration induces expression of the IEG FOS (green) and CENP-A (red) in the adult mouse DG. Left panels show overview of the DG, boxed areas are shown in high power views (right panels). Nuclei were counterstained with DAPI (white). Scale bars represent 100 µm and 50 µm. D, Percentage of dentate gyrus neurons displaying non-centromeric CENP-A, in FOS-negative/ARC-negative versus FOS-positive/ARC-positive cells (the latter denoting nascently activated neurons) following novel-environment exposure. Non-centromeric CENP-A was largely restricted to activated (FOS-positive/ ARC-positive) neurons: 98 of 118 activated neurons were non-centromeric-CENP-A-positive (mean 83%; per-mouse 66, 95, 87%), whereas 0 of 118 non-activated (FOS-negative/ARC-negative) neurons were positive (0% in every animal; matched cell numbers, 38, 42, 38 per mouse). N=3 mice; points are per-mouse percentages. Descriptive; no inferential test. E, Percentage of DG neurons positive for FOS or for non-centromeric CENP-A, each marker quantified independently at home cage (HC), 1 h post novel-environment (NE), and 5 h post-NE. Both markers were transiently induced, rising at 1 h and returning toward baseline by 5 h. FOS: HC 0.32%, 1 h 2.46%, 5 h 0.08% (HC vs 1 h P=0.0002; 1 h vs 5 h P=0.0001). Non-centromeric CENP-A: HC 0.17%, 1 h 2.06%, 5 h 0.05% (HC vs 1 h P=0.0006; 1 h vs 5 h P=0.0004). Unpaired two-tailed t-tests (independent animals per timepoint). FOS and non-centromeric CENP-A were quantified separately and are not compared to one another; the panel illustrates their parallel dynamics. N=3 mice per timepoint; points are per-mouse percentages. F, Total nuclear CENP-A fluorescence intensities in FOS-positive versus FOS-negative dentate gyrus neurons following novel-environment exposure. Total nuclear CENP-A was ∼2.2-fold higher in FOS-positive neurons, in all three animals (per-mouse fold change 2.26, 2.06, 2.18). Fluorescence intensities are background-corrected. N=3 mice, ∼10 cells per condition per mouse; points are per-mouse means. Paired two-tailed t-test on per-mouse means, P=0.0024 (**). Scale bars represent 100 µm for the whole dentate gyrus and 50 µm for high power views. (*, P<0.05, ** P<0.01, *** P<0.001). Figure 1-source **data 1**. Western blot quantification. Figure 1-source **data 2.** Non-centromeric CENP-A in activated and non-activated neurons. Figure 1-source **data 3**. FOS and non-centromeric CENP-A time course. Figure 1-source **data 4.** Total nuclear CENP-A in FOS-positive and FOS-negative neurons. Figure 1-source **data 5**. Original and uncropped western blot images.

Given the activity-dependent regulation of both Cenp-a mRNA and CENP-A protein, we next examined CENP-A protein localization using high resolution imaging. In non-activated neurons (FOS and ARC negative), CENP-A displayed centromeric localization both under control conditions (HC) and following exposure to a NE (Figure 1C). In contrast, in NE-activated conditions, we detected an additional pool of CENP-A. FOS positive neurons showed not only centromeric but also non-centromeric, whole nucleus-enriched CENP-A signal (Figure 1C, 2A). The increase in non-centromeric CENP-A signal occurred in nascently activated neurons (both FOS and ARC positive), with levels returning to the baseline in ARC-only positive neurons (Figure 1C-F and Figure 1-figure supplement 1I-K).

### Non-centromeric CENP-A is dynamically regulated during neuronal activation

To determine which pool of CENP-A is driving its non-centromeric localization, we segmented CENP-A signal into centromeric and total nuclear signal, to derive its non-centromeric levels during neuronal activation. Notably, centromeric CENP-A levels remained unchanged in activated and non-activated neurons, indicating that neuronal activation selectively increases the non-centromeric pool of CENP-A (Figure 2A-D). To directly test localization-dependent CENP-A dynamics, we chronically depleted CENP-A *in vivo* by stereotaxically delivering adeno-associated viruses (AAVs) expressing shRNAs against the endogenous Cenp-a mRNA (Figure 2E-G, Figure 2-figure supplement 1A-B) in postmitotic neurons and analyzed centromeric versus non-centromeric CENP-A levels. Consistent with the previously reported hyperstability of centromeric CENP-A^36, 37^, centromeric CENP-A levels in DG neurons remained unchanged four weeks after viral injection (Figure 2E, F). In contrast, CENP-A knockdown significantly reduced the frequency of neurons exhibiting whole nucleus-enriched CENP-A localization, indicating a selective effect on the activity-induced, non-centromeric CENP-A pool (Figure 2E, G). Live imaging of mouse neural stem cell (mNSC)-derived neurons stably expressing CENP-A-EGFP transgene^38, 39^ further confirmed the hyperstability of centromeric CENP-A and its insensitivity to neuronal activation (Figure 2H, I and Figure 2-figure supplement 1C-D and Figure 2-video 1 and Figure 2-video 2). In contrast, and corroborating the *in vivo* results, the non-centromeric, whole-nuclear pool of CENP-A showed dynamic regulation concomitant with neuronal activity in cultured neurons (Figure 2H-I and Figure 2-figure supplement 1C-D). Together, these data demonstrate that CENP-A exhibits dual localization in activated DG neurons: a stable centromeric pool and a dynamically regulated, non-centromeric fraction that is selectively increased upon neuronal activation.

**Figure 2.**
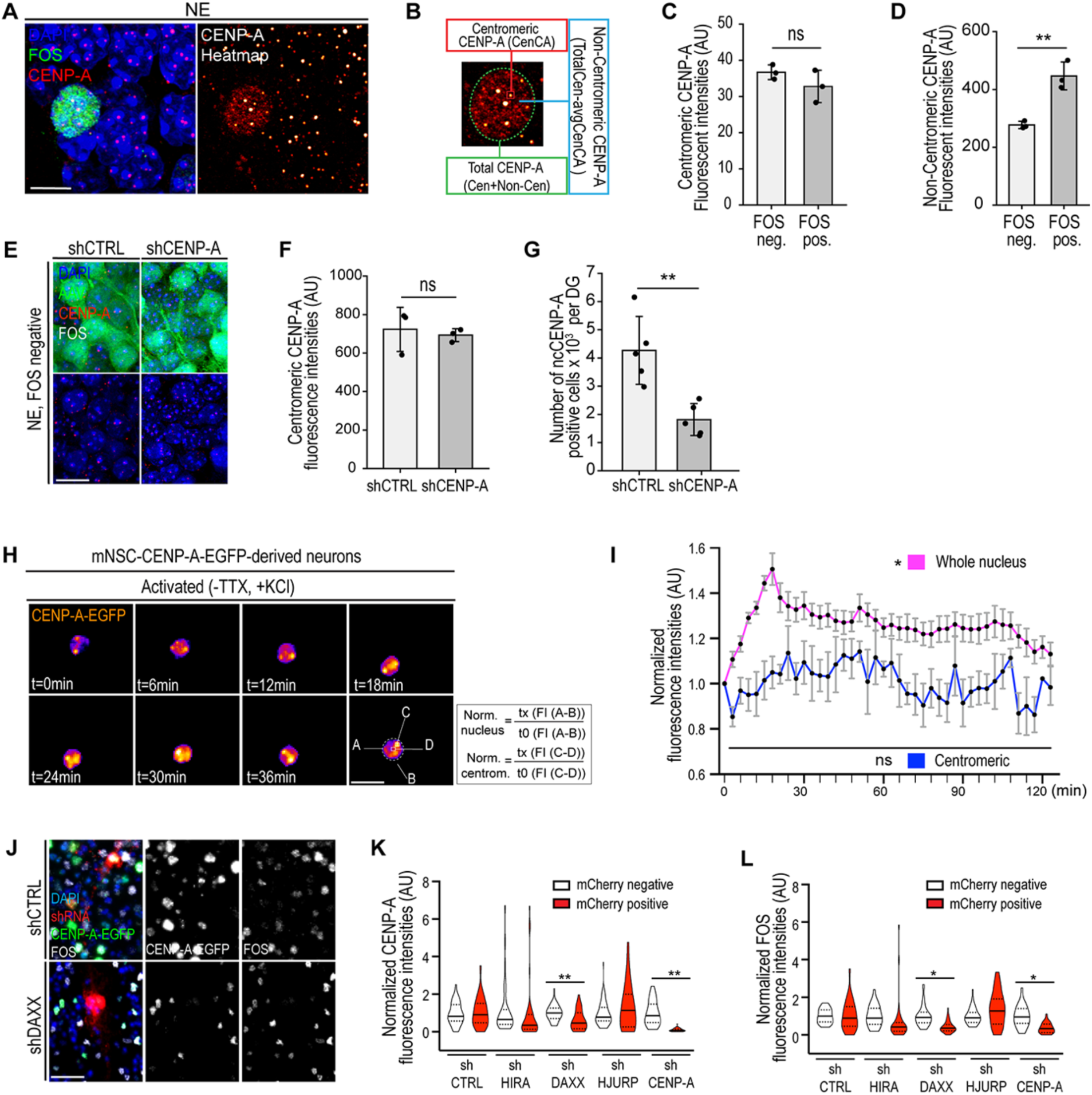
Non-centromeric CENP-A is regulated with neuronal activity in a DAXX-dependent manner. A, Shown is a FOS (green) and CENP-A (red) expressing neuron 1h after exploring a novel environment (NE). Nuclei were counterstained with DAPI (left). Conversion of CENP-A fluorescence intensities into a heat-map (right), highlighting non-centromeric (ncCENP-A) pool present in FOS neurons. Scale bar denotes 5 µm. B, Schematics showing the methodology used to quantify two different pools of CENP-A present in activated neurons. Note that the red-boxed centromeric signals represent examples of several areas with clustered centromeric CENP-A signal. C, Centromeric CENP-A fluorescence intensities in FOS-positive versus FOS-negative dentate gyrus neurons following novel-environment exposure. Centromeric CENP-A did not differ significantly with activation status. N=3 mice, ∼40 cells per condition per mouse (5 centromeres averaged per cell); points are per-mouse means. Paired two-tailed t-test, P=0.21 (ns). D, Non-centromeric CENP-A fluorescence intensities (total nuclear minus centromeric, per cell, averaged per mouse) in FOS-positive versus FOS-negative dentate gyrus neurons. Non-centromeric CENP-A was significantly higher in FOS-positive neurons (1.5- to 1.7-fold), in all three animals. N=3 mice, ∼40 cells per condition per mouse; points are per-mouse means. Paired two-tailed t-test, P=0.009 (**). E, Chronic (4-week) AAV-mediated delivery of either shCTRL or shCENP-A (green) in dentate gyri, followed by the exposure to NE. Scale bar denotes 50 µm. F, Centromeric CENP-A fluorescence intensities in FOS-negative dentate gyrus neurons from mice injected with shCTRL or shCENP-A following novel-environment exposure. Centromeric CENP-A did not differ significantly between conditions, indicating the knockdown spares the centromeric pool. N=3 matched pairs (mice; one shCTRL and one shCENP-A per batch, processed together), ∼60 cells per mouse; paired two-tailed t-test, P=0.59 (ns). G, Total number of non-centromeric CENP-A-positive cells per dentate gyrus in mice injected with shCTRL or shCENP-A following novel-environment exposure (DG neurons scored irrespective of FOS status). Each batch contained one shCTRL and one shCENP-A mouse processed together; data are analysed as matched pairs. shCENP-A reduced the non-centromeric CENP-A-positive population ∼2.35-fold (the knockdown mouse lower than its matched control in every batch). N=5 matched pairs (mice); paired two-tailed t-test, P=0.002 (**). Points are per-animal counts. H, Shown are KCl-stimulated stills of live-imaged CENP-A-EGFP mNSC-derived neurons. Sample was imaged every 3mins, shown is each second frame. Note that the CENP-A-EGFP signal quantification strategy is depicted in the lower right corner. I, Normalized CENP-A-EGFP fluorescence over time in activated mNSC-derived neurons (-TTX, +KCl), whole-nucleus and centromeric signal, each cell normalized to its own first frame. Whole-nucleus CENP-A increased with activation whereas centromeric did not. Lines and markers, mean of 3 independent differentiations; bars, SD across replicates. Activated versus non-activated (Figure 2-figure supplement 1D; unpaired two-tailed Welch’s t-test on per-replicate session means): whole nucleus P=0.02; centromeric P=0.20 (ns). N=3 biological replicates per condition. J, Representative images of mNSC-CENP-A-EGFP-derived neurons treated with the indicated shRNA constructs undergoing KCl-induced activation. Scale bar denotes 5 µm. K, Normalized whole-nucleus CENP-A-EGFP fluorescence in mCherry-positive (shRNA-expressing) versus adjacent mCherry-negative cells, for five shRNA constructs. DAXX and CENP-A knockdown significantly reduced non-centromeric CENP-A in mCherry-positive cells; control, HIRA and HJURP knockdown did not. N=3 biological replicates per construct (∼15 cells per condition per replicate); violins show all cells; the central line indicates the mean of the three per-replicate means. Paired two-tailed t-test on per-replicate means (mCherry+ vs mCherry-), Holm-corrected across the 5 constructs. shDAXX P=0.005 (**); shCENP-A P=0.005 (**); shCTRL, shHIRA, shHJURP ns. L, Normalized FOS fluorescence in mCherry-positive versus adjacent mCherry-negative cells, for the same five shRNA constructs. DAXX and CENP-A knockdown significantly reduced FOS in mCherry-positive cells; control, HIRA and HJURP knockdown did not (after correction for multiple comparisons). N=3 biological replicates per construct (∼15 cells per condition per replicate); violins show all cells; the central line indicates the mean of the three per-replicate means. Paired two-tailed t-test on per-replicate means, Holm-corrected across the 5 constructs. shDAXX P=0.030; shCENP-A P=0.023; shCTRL, shHIRA, shHJURP ns (shHIRA raw P=0.063, not significant after Holm correction). (*, P<0.05, ** P<0.01, *** P<0.001). Figure 2-source **data 1**. Centromeric CENP-A, cell-level data. Figure 2-source **data 2.** Non-centromeric CENP-A, cell-level data. Figure 2-source **data 3**. Centromeric CENP-A after knockdown, cell-level data. Figure 2**-source data 4.** Non-centromeric-CENP-A-positive cells after knockdown. Figure 2**-source data 5.** Live imaging, activated and non-activated, per-cell traces. Figure 2**-source data 6.** Chaperone screen, CENP-A, cell-level data. Figure 2-source **data 7.** Chaperone screen, FOS, cell-level data.

### DAXX is necessary for activity-induced accumulation of non-centromeric CENP-A

To gain further insights into the mechanism underlying activity-dependent accumulation of non-centromeric CENP-A, we used a candidate-based shRNA knockdown approach to identify histone chaperones that may contribute to this process. We assayed for the presence and extent of non-centromeric CENP-A signal, using both the CENP-A-EGFP transgene and direct immunostaining of endogenous CENP-A, in mNSC-derived neurons expressing shRNAs against three histone chaperones previously associated with CENP-A deposition (Figure 2J-L and Figure 2-figure supplement 1E-H)^7, 40, 41^. As non-centromeric localization of CENP-A has been previously linked to H3.3 chaperones in the context of proliferating cancer cells with massive overexpression of CENP-A^40, 42^, we probed the contribution of HIRA and DAXX, along with the CENP-A-specific chaperone HJURP, in driving activity-dependent deposition of CENP-A. Strikingly, depletion of DAXX, but not of HIRA or HJURP, led to a significant decrease in non-centromeric CENP-A deposition (Figure 2K and Figure 2-figure supplement 1F-G). This effect was observed both with the CENP-A-EGFP transgene (Figure 2K) and with direct immunostaining of endogenous CENP-A (Figure 2-figure supplement 1F-G), arguing against a transgene-related artifact. Consistent with a functional role for this DAXX-dependent CENP-A pool, DAXX depletion also reduced the activity-induced induction of FOS (Figure 2L and Figure 2-figure supplement 1H). Thus, our results identify a pathway contributing to activity-dependent accumulation of non-centromeric CENP-A under physiological conditions in postmitotic neuronal cells.

### Non-centromeric CENP-A associates with transcriptionally competent chromatin in neurons

To characterize the genomic occupancy of non-centromeric CENP-A, we performed *in vivo* ChIP-seq on DG neurons^43^. As expected, a large fraction of CENP-A reads mapped to minor satellite repeats corresponding to annotated murine centromeres; in addition, we detected a distinct non-centromeric fraction of CENP-A (Figure 3-figure supplement 1A). This non-centromeric fraction preferentially localized to promoter regions, defined as ±1kb relative to transcription start sites (TSS) (Figure 3A), and to a lesser extent to gene bodies (Figure 3-figure supplement 1D). In contrast, non-neuronal hepatocytes do not exhibit comparable promoter localization of CENP-A^43^, indicating that promoter association of non-centromeric CENP-A may be a neuron-specific property (Figure 3A). Unsupervised clustering of CENP-A-occupied promoters identified four distinct genomic clusters (Figure 3A). Integration with independent neuron-specific epigenomic datasets (for GEO access numbers refer to Materials and Methods) revealed that these clusters were associated with features of open chromatin, including CpG islands and enrichment of H3K27ac (Figure 3A). Consistent with their localization within transcriptionally active chromatin, CENP-A-enriched promoter clusters also showed strong correspondence with H3K4me3, RNA Polymerase II, CREB-binding protein (CBP), and chromatin accessibility measured by ATAC-seq (Figure 3B). Similarly, unsupervised clustering of CENP-A-occupied gene bodies identified three clusters (Figure 3-figure supplement 1D), associated with CpG islands, H3K27ac, H3K4me3, RNA Polymerase II, and accessible chromatin, and corresponding to genes transcriptionally induced upon neuronal activation (Figure 3-figure supplement 1E)^17^.

**Figure 3.**
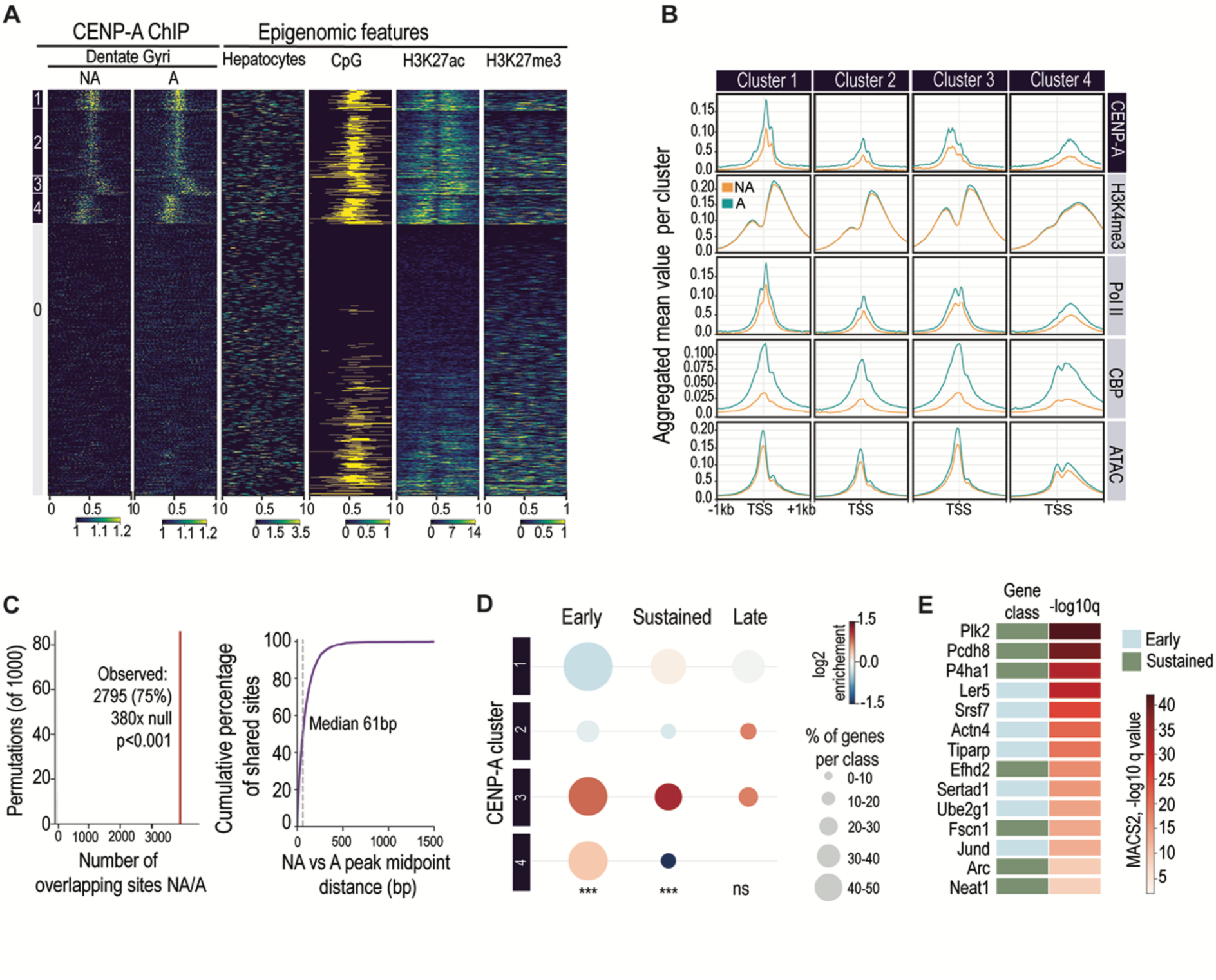
CENP-A occupancy maps to activity-gene promoters in Dentate Gyrus neurons. A, Genome-wide CENP-A occupancy in DG neurons in non-activated (NA) and activated (A) states, displayed as a heatmap over CENP-A-occupied promoter regions grouped into four clusters (plus a low-confidence background group). Adjacent columns show reused epigenomic features for context: hepatocyte CENP-A (non-neuronal specificity control), CpG islands, H3K27ac and H3K27me3. NA and A are shown to illustrate conserved occupancy (quantified in C). Clustering was performed via k-means clustering. B, Aggregated mean signal per promoter cluster for CENP-A and reused epigenomic marks (H3K4me3, Pol II, CBP, ATAC), in non-activated (orange) and activated (teal) states, across the four promoter clusters. NA and A shown to demonstrate conserved occupancy. The NA/A traces for the epigenomic marks are independent published datasets; divergence in those rows (notably CBP) reflects the external datasets, not a CENP-A activity effect. C, CENP-A occupancy is conserved between non-activated (NA) and activated (A) states. Left, the number of activated CENP-A peaks overlapping non-activated peaks (observed, red) versus a within-chromosome permutation null (1,000 shuffles); 2,795 of 3,733 activated peaks (75%) overlap non-activated peaks, ∼380-fold above the null expectation (P<0.001). Right, cumulative distribution of midpoint distances between overlapping peaks (median 61 bp; 99% within 500 bp), indicating the two states occupy the same genomic positions. D, Distribution of activity-induced gene temporal classes (early, sustained, late; based on^32^) across the four CENP-A promoter clusters. Dot size, percentage of each temporal class in each cluster; colour, log2 enrichment over the cluster-size background (blue, depletion; red, enrichment). Early and sustained activity-induced genes were significantly enriched in cluster 3 (early 1.79-fold, P=3.8e-5; sustained 2.21-fold, P=6.5e-4, chi-square versus cluster-size proportions), whereas late-response genes were not (P=0.10); housekeeping genes showed no significant clustering (P=0.48). E, Selected activity-induced genes (early or sustained) whose promoter CENP-A peak was detected reproducibly in cluster 3. Left column, temporal class; right column, peak occupancy confidence (MACS2 −log10 q, maximum across the two libraries). Colour reflects statistical confidence of occupancy, not CENP-A abundance. Figure 3-source **data 1.** Promoter cluster assignment and gene lists. Figure 3-source **data 2.** Peak overlap and permutation null. Figure 3-source **data 3.** Temporal class enrichment. Figure 3-source **data 4.** Gene occupancy confidence.

Because IEGs are induced with distinct temporal kinetics, we asked whether CENP-A occupancy was preferentially associated with particular classes of activity-induced genes. Among the four promoter clusters, cluster 3 was significantly enriched for early and sustained activity-induced genes (1.79-fold, P=3.8e-5 and 2.21-fold, P=6.5e-4, respectively), but not for late-response or housekeeping genes (Figure 3D). Individual early and sustained activity-induced genes with reproducible promoter CENP-A occupancy included canonical IEGs such as JunD and Arc (Figure 3E). Thus, non-centromeric CENP-A occupies the promoters of a defined set of activity-induced genes.

Finally, we compared CENP-A occupancy between non-activated and activated neurons. CENP-A occupancy was highly conserved between the two states: 75% of activated CENP-A peaks overlapped with non-activated peaks (∼380-fold above a permutation null, P<0.001), and overlapping peaks occupied nearly identical genomic positions (median offset 61 bp; Figure 3C). This indicates that non-centromeric CENP-A is stably pre-positioned at activity-dependent gene promoters under basal conditions, rather than being recruited *de novo* upon activation. Together, these findings reveal that non-centromeric CENP-A in neurons is constitutively poised at transcriptionally competent, activity-associated genomic regions, providing a chromatin context for the activity-dependent increase in the non-centromeric CENP-A pool.

### Activity-regulated CENP-A is necessary for FOS and ARC induction

Given the activity-dependent regulation of CENP-A, we next probed for the functional relevance of CENP-A upregulation. To this end, we downregulated CENP-A in DG neurons by injecting AAVs expressing shRNAs directed against the coding mRNA of CENP-A (Figure 4A) and assayed for the presence of two canonical IEGs, FOS and ARC^44, 45^. Virus-mediated downregulation of CENP-A caused a substantial decrease in the number of FOS-labeled cells in the DG after NE exposure, as quantified by two complementary approaches: immunohistochemical tissue analysis and fluorescence activated cell sorting (FACS) (Figure 4B-C). In analogy to FOS-labeled cells, the number of ARC-positive cells was reduced after NE exploration upon CENP-A knockdown (Figure 4-figure supplement 1A-B). Notably, depletion of Fos did not affect activity-dependent Cenp-a expression (Figure 4-figure supplement 1C)^33^. To further define the temporal relationship between CENP-A accumulation and IEG induction, we performed imaging with high temporal resolution of CENP-A and FOS dynamics in mNSC-derived neurons following neuronal activation. Whole-nuclear accumulation of CENP-A preceded the induction of FOS (Figure 4-figure supplement 1D-E), suggesting that non-centromeric CENP-A accumulation precedes IEG induction following neuronal activation. Consistent with this model, depletion of DAXX in mNSC-derived neurons significantly reduced both the frequency and intensity of FOS induction (Figure 2L and Figure 2-figure supplement 1H), corroborating the requirement for non-centromeric CENP-A accumulation for activity-mediated IEG expression. These data show that downregulation of CENP-A results in decreased capacity of hippocampal neurons to respond to external stimuli, as evidenced by the reduced induction of two key markers of neuronal activity, FOS and ARC. Thus, CENP-A upregulation and its whole nucleus enrichment are tightly connected to neuronal activation.

**Figure 4.**
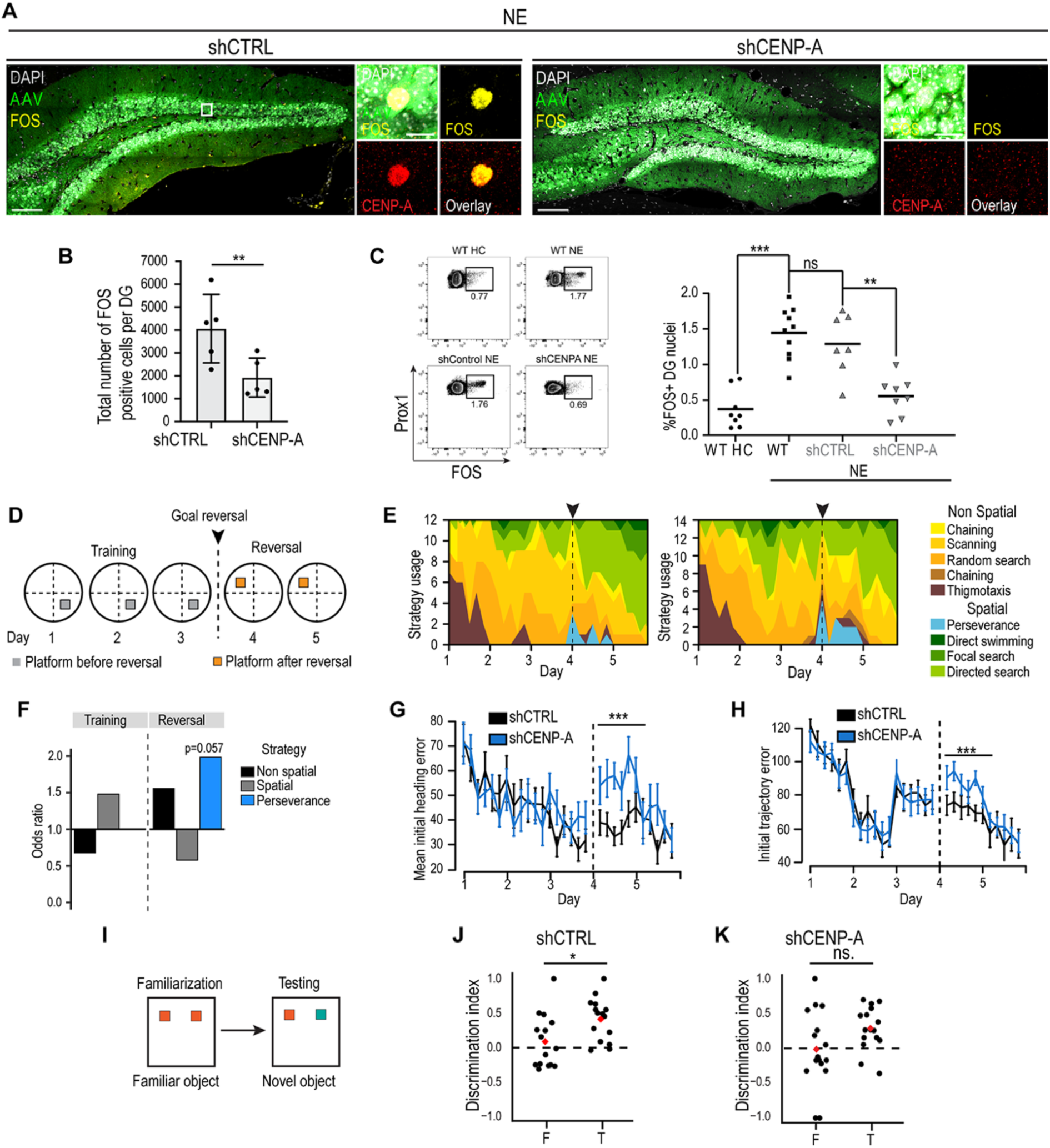
CENP-A knockdown impairs the activity-induced response in vivo. A, AAV-mediated knockdown of CENP-A (shCTRL, left panel; shCENP-A, right panel; green) reduces CENP-A levels (red) and number of FOS-labeled neurons (yellow) after NE. Boxed areas are shown in high power views and single channels. Overview images consist of individual images that were electronically stitched together. Scale bars represent 100 µm. B, Number of FOS-positive cells per dentate gyrus in shCTRL- and shCENP-A-injected mice following novel-environment exposure. Each batch contained one shCTRL and one shCENP-A mouse, injected, and 4 weeks later exposed to NE and perfused together (randomized block design); data are analysed as matched pairs. shCENP-A reduced the number of FOS-positive cells by ∼53% (2.12-fold), with the knockdown mouse lower than its matched control in every batch. N=5 matched pairs (mice); paired two-tailed t-test, P=0.0049 (**). Points are per-animal counts. C, Percentage of FOS-positive dentate gyrus nuclei determined by FACS. Novel-environment exposure induced FOS in non-injected mice (home cage 0.37% vs novel environment 1.45%; P=2.8e-6), confirming the paradigm. Control hairpin injection did not alter the response (non-injected NE vs shCTRL NE; P=0.43). shCENP-A reduced the FOS response 2.32-fold relative to shCTRL (1.29% vs 0.56%; P=0.0014). N=8 (HC), 10 (NE), 7 (shCTRL), 8 (shCENP-A) mice; unpaired two-tailed t-tests. Points are per-mouse percentages. D, Schematic of the Morris water maze paradigm that includes reversal learning of a novel platform position starting on day 4 of the task. E, Visualization of search strategies during acquisition (day 1-3) and reversal learning (day 4-5, indicated by dotted line). F, Odds ratio of strategy usage over learning days in control mice and after CENP-A knockdown (p values were determined by fitting a generalized linear model with binomial distribution). G, Initial heading error over the learning period. Dotted line indicates start of reversal learning. ***, P<0.001 on day 4 (unpaired two-tailed t-test on data binned by day). H, Initial trajectory error over the learning period. Dotted line indicates start of reversal learning. ***, P<0.001 on day 4 (unpaired two-tailed t-test on data binned by day). I, Schematic of the novel object recognition (NORT) test paradigm. J-K, Discrimination index during familiarization (F) and test (T), when a familiar and a novel object were presented, in shCTRL (J) and shCENP-A (K) mice, computed over the first 30 s of object exploration. Control mice showed a significant preference for the novel object at test (P=0.017), whereas the increase in shCENP-A mice did not reach significance (P=0.066, ns). N=15 (shCTRL) and 16 (shCENP-A) mice; two-sample two-tailed t-tests comparing phases within each group. Points are individual mice; red diamonds show group means. Figure 4-source **data 1.** FOS-positive cells after knockdown. Figure 4-source **data 2.** FACS quantification. Figure 4-source **data 3.** Morris water maze raw tracking data (deposited at Zenodo, DOI: 10.5281/zenodo.23327295). Figure 4-source **data 4**. Novel object recognition, discrimination index.

### CENP-A is necessary for hippocampus-dependent learning and memory

We next tested if hippocampus-dependent learning and memory depends on the activity-regulated expression of CENP-A by analyzing mice in hippocampus-dependent behavioral tasks after CENP-A knockdown. Again, we downregulated CENP-A in the DG by injecting AAVs expressing shRNA against CENP-A. Four weeks after viral injection, we first probed for a CENP-A function in hippocampus-dependent behavior, by analyzing mice in a spatial memory task of the Morris water maze (Figure 4D). Both control mice and mice with CENP-A knockdown learned the behavioral task (Figure 4-figure supplement 1F-H); however, CENP-A knockdown mice showed a trend (p = 0.057) of an altered, hippocampus-dependent search strategy after platform location change on day 4 and 5 of the task (Figure 4E-F). Notably, knockdown of CENP-A caused impaired learning parameters compared to controls, exhibiting enhanced heading and trajectory errors when re-learning the novel position of the escape platform (Figure 4G-H)^46^. To extend the water maze-based findings, mice were analyzed in an additional behavioral task, using a hippocampus-dependent version of the novel object recognition task (NORT) (Figure 4I). Control mice showed a significant preference for the novel object at test, whereas the preference of CENP-A knockdown mice did not reach significance (Figure 4J-K), suggesting impaired hippocampus-dependent object recognition after CENP-A knockdown. Thus, our experiments reveal that CENP-A shows activity-dependent regulation, influences IEGs (FOS and ARC), and contributes to hippocampus-dependent learning and memory in mice.

### Activity-mediated neuronal function of CENP-A is conserved between mice and humans

Next, we asked if the unexpected non-centromeric function of CENP-A in the context of neuronal activation is conserved in human neural tissues. We used human embryonic stem cell (hESC)-derived forebrain organoids, allowing for the analyses of activity-dependent gene expression in self-organized, human neural networks^47–50^.

After maturing for more than 90 days, regionalized forebrain organoids were transduced with AAVs expressing Channelrhodopsin-2 (ChR2) driven by the regulatory elements of the human synapsin-1 promoter (hSyn1)^51^. Two weeks later organoids were exposed to blue light, thereby optogenetically activating targeted neurons (Figure 5A and Figure 5-figure supplement 1A). Optogenetic activation of neurons within organoids drove FOS expression in CTIP2-expressing neurons but not in SOX2-labeled progenitor cells (Figure 5A-C). Furthermore, optogenetic stimulation of human neurons caused a robust increase of whole-nucleus enriched CENP-A in FOS positive cells (Figure 5D-F). In contrast, the levels of centromeric CENP-A remained unaffected in ChR2-activated neurons (Figure 5E), corroborating the mouse in vivo results. Finally, we analyzed if - in analogy to the mouse brain - CENP-A is also required for IEG presence in human neuronal networks. AAV-mediated knockdown of human CENP-A substantially reduced the number of FOS-labeled neurons upon optogenetic stimulation compared to control neurons (Figure 5G-H), an effect that was not due to differences in transduction efficiency between the different experimental conditions (Figure 5I). Importantly, ChR2-positive hairpin-negative cells activated normally, confirming that the effect is specific to CENP-A knockdown (Figure 5H). Collectively, our data indicate a conserved regulation and function of CENP-A between mouse and human neurons in the context of synaptic activation.

**Figure 5.**
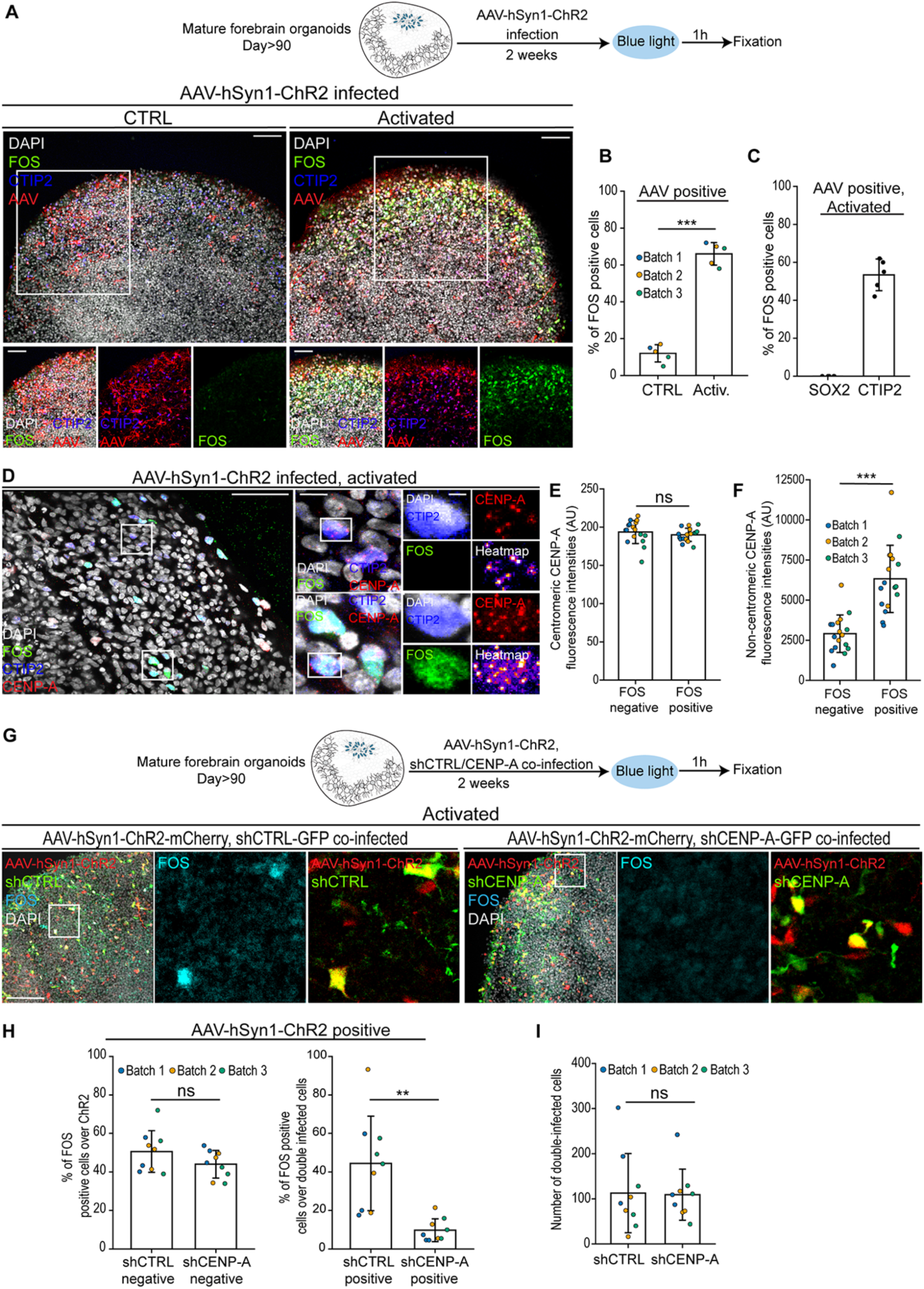
Activity-mediated function of CENP-A is conserved between mice and humans. All panels use AAV-hSyn1-ChR2-transduced forebrain organoid neurons. “CTRL” denotes AAV-infected cells not exposed to blue light; “Activated” denotes AAV-infected cells exposed to blue light. The biological replicate is the cortical unit; the differentiation batch is accounted for (see Methods). A, Schematic of experimental approach consisting of infection of mature human organoids with AAVs expressing Channelrhodopsin-2 (ChR2) under the control of the human Synapsin1 (hSyn1) promoter. Left panels show representative examples of organoids infected with AAV-hSyn1-ChR2 (red) under control conditions (left) or after exposure to blue light (right) expressing FOS (red) in CTIP2 (blue)-labeled neurons. Nuclei were counterstained with DAPI (white). Scale bars represent 50 µm. B, Percentage of AAV-infected (AAV-positive) cells that are FOS-positive, without (CTRL) and with (Activated) blue-light stimulation. Blue-light activation increased the FOS-positive fraction ∼5.5-fold (12.0% vs 66.0%), consistently across all three batches. N=5 cortical units per condition from 3 differentiation batches; unpaired two-tailed Welch’s t-test, P<0.0001. Points are per-cortical-unit percentages. C, Percentage of FOS-positive cells that are SOX2-positive (progenitors) or CTIP2-positive (neurons) in activated organoids. FOS-positive cells were essentially all neurons: 0% were SOX2-positive progenitors (across all organoids), whereas a mean of 53% were CTIP2-positive neurons. N=5 organoids from 3 differentiation batches (∼50 cells scored per organoid); descriptive. Points are per-organoid percentages. D, Representative examples of organoids infected with AAV-hSyn1-ChR2 (unstained) under activated conditions (exposure to blue light) (right) expressing CENP-A (red) in FOS-(green), CTIP2 (blue)-labeled neurons. Boxed areas are shown in high power views. Right lower panels show CENP-A signal converted to heat-map. Nuclei were counterstained with DAPI (white). Scale bars represent 50 µm, 10 µm, 2 µm, respectively. E, Centromeric CENP-A fluorescence intensities in FOS-positive versus FOS-negative neurons within activated (blue-light-stimulated) organoids. Centromeric CENP-A did not differ significantly with activation status. N=17 cortical units from 3 differentiation batches; paired comparison at the cortical-unit level (batch accounted for), P=0.39 (ns). Points are per-cortical-unit means. F, Non-centromeric CENP-A fluorescence intensities (total nuclear minus centromeric) in FOS-positive versus FOS-negative neurons within activated organoids. Non-centromeric CENP-A was significantly higher in FOS-positive neurons (2.18-fold), an increase observed in all 17 cortical units. N=17 cortical units from 3 differentiation batches; paired comparison at the cortical-unit level (batch accounted for), P<0.0001. Points are per-cortical-unit means. G, Schematic of experimental approach consisting of co-infection of mature human organoids with AAV-hSyn1-ChR2 and AAVs expressing control shRNA or shRNA targeting human CENP-A. Lower panels show representative examples of organoids infected with AAV-hSyn1-ChR2 (red) and shCTRL (green; left 3 panels) expressing FOS (blue) after light-induced activation. Three panels on the right show loss of FOS expression (blue) after co-infecting cells with AAV-hSyn1-ChR2 and AAV expressing shCENP-A (green). Boxed areas are shown in high power views. Nuclei were counterstained with DAPI (white). Scale bar represents 50 µm. H, Left, percentage of FOS-positive cells among ChR2-positive, hairpin-negative cells, in shCTRL versus shCENP-A conditions. Cells that received the activating ChR2 but not the knockdown hairpin activated equivalently between conditions, confirming matched activation capacity independent of the hairpin (50.6% vs 44.0%; P=0.15, ns). Right, percentage of FOS-positive cells among double-infected (ChR2-positive and hairpin-positive) cells. In cells receiving both the activating ChR2 and the knockdown hairpin, shCENP-A reduced FOS 4.6-fold (44.4% vs 9.7%), consistently across all three batches (P=0.003). N=9 cortical units from 3 differentiation batches; unpaired two-tailed Welch’s t-tests at the cortical-unit level. Points are per-cortical-unit percentages. I, Number of double-infected (ChR2-positive and hairpin-positive) cells per cortical unit in shCTRL versus shCENP-A conditions. Infection efficiency did not differ between conditions, confirming that the reduced FOS response in shCENP-A double-infected cells (H) is not attributable to differential infection (112.6 vs 109.1; P=0.91, ns). N=9 cortical units from 3 differentiation batches; comparison at the cortical-unit level (batch accounted for). Points are per-cortical-unit counts. (*, P<0.05, ** P<0.01, *** P<0.001). Figure 5**-source data 1.** FOS induction in organoids. Figure 5-source **data 2.** SOX2 and CTIP2 identity of FOS-positive cells. Figure 5-source **data 3.** Centromeric CENP-A in organoids. Figure 5-source **data 4.** Non-centromeric CENP-A in organoids. Figure 5**-source data 5.** FOS after CENP-A knockdown in organoids. Figure 5**-source data 6**. Double-infected cell counts.

## DISCUSSION

CENP-A has been extensively studied for its essential role in specifying centromere identity and ensuring faithful chromosome segregation during cell division^1, 52^. In contrast, a functional role for non-centromeric CENP-A in post-mitotic cells under physiological conditions has not been described. Here, we identify a co-option of CENP-A function in hippocampal neurons, where it contributes to activity-dependent gene expression and supports hippocampus-dependent learning and memory. Unlike its canonical mitotic role, neuronal activation induces accumulation of a non-centromeric nuclear pool of CENP-A that influences the efficiency of activity-dependent neuronal responses. Notably, despite rapid sequence divergence of CENP-A across species, activity-dependent regulation of CENP-A in neurons is conserved between mice and humans, underscoring the functional significance of CENP-A repurposing.

Previous studies in cancer cell lines have reported non-centromeric CENP-A accumulation in the context of pathological CENP-A overexpression^40, 42^. Extending these observations, we demonstrate that under physiological conditions in post-mitotic neurons, neuronal activation induces a distinct non-centromeric pool of CENP-A that contributes to activity-dependent gene expression programs. Furthermore, ChIP-seq analysis reveals that this non-centromeric pool is poised at transcriptionally competent genomic regions under basal conditions, suggesting a chromatin architecture primed for rapid transcriptional responses. While these ChIP-seq analyses are based on a limited number of biological replicates, the genomic distribution and cluster identity of non-centromeric CENP-A reported here are consistent between conditions and supported by cross-correlation with independent published datasets. Activity-dependent accumulation of non-centromeric CENP-A depends on the histone chaperone DAXX and is consistent with the formation of heterotypic nucleosomes comprising CENP-A and canonical histone H3.3, as previously described in proliferating cells^40^.

A key distinction between centromeric and non-centromeric CENP-A in neurons lies in their stability. While centromeric CENP-A is hyperstable and insensitive to neuronal activity, the non-centromeric pool is highly dynamic and responds rapidly to changes in neuronal activity. This dynamic behaviour relies on nascent CENP-A expression. The low levels of CENP-A detected outside centromeres under basal conditions may reflect the intrinsic nucleosomal stability conferred by CENP-A itself, as incorporation of CENP-A outside centromeres has been shown to stabilize its binding partner H4 ^37^. In contrast, the hyperstability of centromeric CENP-A depends on interactions with CCAN components such as CENP-B, CENP-C, and CENP-N^36, 53, 54^, factors that are limiting in post-mitotic neurons. We propose that the balance between the intrinsic stability of CENP-A-containing nucleosomes and limiting CCAN availability creates a permissive chromatin environment that enables activity-dependent accumulation of non-centromeric CENP-A. In this context, non-centromeric CENP-A may act as a priming factor that facilitates transcriptional competence during neuronal activation. Structurally, the shorter αN helix of CENP-A compared to canonical H3 reduces DNA contacts at nucleosome entry and exit sites, increasing DNA flexibility^55–57^. This property raises the possibility that CENP-A incorporation promotes a dynamic chromatin state permissive to transcription factor binding^58^. Future work will be required to identify interaction partners and post-translational modifications of non-centromeric CENP-A in neurons, which may further modulate activity-dependent transcription.

Importantly, our findings resonate with recent work showing that ectopic non-centromeric CENP-A accumulation in human cancer cells can influence transcriptional programs associated with epithelial-mesenchymal plasticity^59^. While this study focused on pathological CENP-A overexpression in proliferating cells, our results reveal a conceptually analogous mechanism operating under physiological conditions in post-mitotic neurons, where activity-dependent CENP-A accumulation contributes to activity-dependent gene expression without compromising centromere integrity. Together, these studies point to a broader principle whereby non-centromeric CENP-A may act as a reversible epigenetic modulator of cell-state plasticity across diverse biological contexts.

Despite the rapid evolution of centromeric DNA and CENP-A protein sequences, it is tempting to speculate that the conserved co-option of a centromeric histone variant for activity-dependent transcription enables the translation of experience into long-lasting changes in neuronal function. Indeed, IEG expression has been observed across species with highly divergent centromeric architectures, from mammals to invertebrates^60, 61^. Determining when during evolution CENP-A acquired a role in learning and memory remains an open question. Nonetheless, our work reveals that in mammalian neurons, repurposing CENP-A outside of mitosis provides a powerful mechanism to regulate activity-dependent gene expression, positioning CENP-A as a novel chromatin component underlying neuronal plasticity and learning.

## MATERIALS AND METHODS

### Animal husbandry

Animal testing was sanctioned by the veterinary authorities of Zurich, Switzerland, or by the Landesdirektion Sachsen, adhering to both European and national guidelines. Mice were collectively accommodated in ventilated cages, following a 12-hour dark/light regimen, and provided unrestricted access to food and water in an environment free from pathogens. Where applicable, C57BL/6 of mixed sex were used.

### Mouse neural stem cell cultures

Mouse neural stem cells (mNSCs) were derived from the hippocampus of adult C57BL/6 animals^62^. Isolated cells were cultured in DMEM/F12 GlutaMax medium supplemented with N2, B27, antibiotics, EGF (20 ng/ml, Thermo Fisher Scientific), FGF-2 (20 ng/ml, PeproTech) and Heparin (5 mg/ml, Sigma-Aldrich). Plasmids were introduced using the AMAXA electroporation system (Lonza) ^63^. mNSCs were differentiated into neurons by growth factor withdrawal (no addition of EGF and FGF). Cells were grown under standard conditions (37 °C, 5% CO2).

### Forebrain-regionalized cerebral organoids

H9-hESCs were cultured in standard 37 °C and 5% CO2 conditions in feeder-free conditions on hESC-grade Matrigel (Corning) coated plates^48, 64^. Cells were maintained in mTeSR Plus (Stem Cell Technologies) in the absence of antibiotics and media was changed every two days. Forebrain-regionalized cerebral organoids were produced using the previously published protocol with some modifications^64, 65^. In brief, half a million hESCs were passaged into an AggreWell-800 well (24-well plate, Stem Cell Technologies) pretreated with Anti-Adherence Rinsing Solution (Stem Cell Technologies). Cells were kept 24 h in mTeSR Plus with 10 µM Y-27632 until they formed embryoid bodies (EBs). EBs were collected the next day (day 1) and transferred to Ultra-Low Attachment Plates (Sigma-Aldrich) with TeSR-E5 (Stem Cell Technologies) supplemented with 2 µM Dorsomorphin (Sigma-Aldrich) and 2 µM A83-01 (Tocris) and fed on day 3. On day 4 and day 5, EBs were adapted to Induction media (TeSR-E5 supplemented with 1 µM CHIR99021, Stem Cell Technologies; and 1 µM SB431542, Stem Cell Technologies) performing half-volume media changes. On day 7 EBs were embedded in growth factor reduced Matrigel (Corning) to induce neural tube formation. From day 7 to day 14 organoids were fed with induction media every two days. Finally, organoids were removed from the Matrigel and placed in miniaturized spinning bioreactors at 100 rpm and maintained in Differentiation media (DMEM-F12 supplemented with N2, B27, penicillin, streptomycin, NEAA, 100 µM β-mercaptoethanol, and 2.5 µg/mL insulin). At day 50, growth media was switched to Maturation media (Neurobasal (Gibco), 1X B27 Supplement, 1X Penicillin/Streptomycin, 1X 2-Mercaptoethanol, 0.2 mM Ascorbic Acid, 20 ng/ml BDNF (PeproTech), 20 ng/ml GDNF (PeproTech) and 0.5 mM cAMP (Sigma)), to promote and enhance neuronal survival and synaptic connectivity. Organoids used in experiments were at least 90 days old.

### Immunofluorescence

Cultured cells and forebrain organoids were fixed with 4% PFA and incubated for 10 min or 30 min, respectively ^64^. Brains were collected after transcardiac perfusion, kept in 20% sucrose overnight, followed by cutting into 40 µm sections using a sliding microtome^62^. Antibodies used were: anti-mCherry (1:250, rabbit, Thermo Fisher Scientific), anti-Fos (1:500, goat sc-52, Santa Cruz, discontinued), anti-Fos (1:500, rabbit, CST 9F6), anti-Arc (1:500, guinea pig, Synaptic Systems 156004), anti-CTIP2 (1:200, rat, Abcam 18465), anti-SOX2 (1:200, goat, Santa Cruz 17320), and anti-CENP-A (1:250, rabbit, CST 2048), followed by incubation with fluorescently labeled secondary antibodies. Nuclei were counterstained with DAPI (1:1000, Sigma).

### Western blotting

Hippocampi from single PTZ-treated and Saline-injected mice were harvested and lysed in RIPA buffer containing 150 mM NaCl, 1% IGEPAL, 0.5% sodium-deoxycholate, 0.1% SDS, 25 mM Tris pH 7.4. Following tissue dissociation, Benzonase nuclease was added to 1% total volume. Following DNA digestion, sodium concentration was adjusted to 300 mM by addition of concentrated NaCl. Chromatin-bound fraction of CENP-A was then purified via Amicon Centrifugal filters (Millipore, UFC910024). Following isolation, samples were subjected to western blotting using 12% precast gels (Bio-Rad) and immunoblotting against CENP-A (CST, 2048) and GAPDH (mouse, HyTest, 5G4). IRDye800CW-coupled anti-rabbit (LI-COR Biosciences) and DyLight680-coupled anti-mouse (Rockland Immunochemicals, Gilbertsville, PA) secondary antibodies were used prior to detection on an Odyssey near-infrared scanner (LI-COR Biosciences, Lincoln, NE). Immunoblot signals were quantified using the Odyssey software and CENP-A signal was normalized to loading control (GAPDH).

### Cloning and shRNA design

Hairpin designed to deplete endogenous mouse CENP-A transcript was designed via the GPP Web portal (Broad Institute, https://portals.broadinstitute.org/gpp/public/), followed by subcloning into pUEG-U6-EF1A-H2B-mCherry. Efficiency of knockdown was assayed in proliferating mNSCs, 72 h post-electroporation. The most efficient hairpin (5’-GCGCAGAAGACAGAAATTCAT-3’) was used as a template for creation of AAV-based vectors, where the expression of hairpin is driven by the truncated human Synapsin promoter and infected cells are visualized via EGFP signal. For depletion of human CENP-A transcript, a previously published CENP-A hairpin (5’-AGGAGATCCGAAAGCTTCA-3’)^66^ together with two additional ones provided by Invitrogen’s BLOCK-iT RNAi Designer (5’-GCAGCAGAAGCATTTCTAGT-3’ and 5’-GAGTTACTCTCTTCCAAAGG-3’) were used as a template to create AAV-based vectors for transduction of forebrain organoids, analogously to the above-mentioned mouse Cenpa-targeting AAV vectors.

### Chromatin immunoprecipitation (ChIP)

CENP-A chromatin immunoprecipitation was performed according to Iwata-Otsubo et al.^43^. Dentate gyri from 4 C57BL/6 wild-type mice per experimental condition were microdissected and flash frozen. Tissue was homogenized in 1 ml ice-cold Buffer 1 (0.32 M sucrose, 60 mM KCl, 15 mM NaCl, 15 mM Tris-Cl, pH 7.5, 5 mM MgCl2, 0.1 mM EGTA, 0.5 mM DTT, 0.1 mM PMSF, 1 mM leupeptin/pepstatin, 1 mM aprotinin) per g of tissue by Dounce homogenization, first with pestle A and then pestle B. Homogenate was strained through a 70 µm cell strainer and spun for 5 min at 1000g, 4 °C. The pellet was then resuspended in 1 ml Buffer 1 containing 0.2% IGEPAL for a final concentration of 0.1%. Samples were incubated for 10 min on ice with occasional vortexing. 2 ml nuclei were gently layered on top of 4 ml ice-cold Buffer 3 (1.2 M sucrose, 60 mM KCl, 15 mM NaCl, 5 mM MgCl2, 0.1 mM EGTA, 15 mM Tris, pH 7.5, 0.5 mM DTT, 0.1 mM PMSF, 1 mM leupeptin/pepstatin, 1 mM aprotinin) and centrifuged at 7684g for 30 min at 4 °C with no brake. Pelleted nuclei were resuspended in 2.5 ml of Wash Buffer A (0.34 M sucrose, 15 mM HEPES, pH 7.4, 15 mM NaCl, 60 mM KCl, 4 mM MgCl2, 1 mM DTT, 0.1 mM PMSF, 1 mM leupeptin/pepstatin, 1 mM aprotinin), flash-frozen in liquid nitrogen, and stored at −80 °C. After thawing in luke-warm water, nuclei were supplemented with 3 mM CaCl2 and digested with 0.1 U/µg of MNase (New England Biolabs). The reaction was stopped with 10 mM EGTA on ice for 5 min, and an equal volume of 2× post-MNase Buffer (40 mM Tris, pH 8, 220 mM NaCl, 4 mM EDTA, 2% Triton X-100, 0.5 mM DTT, 0.5 mM PMSF, 1 mM leupeptin/pepstatin, 1 mM aprotinin) was added. Samples were centrifuged at 14000g for 30 min at 4 °C. After spinning, the samples were pre-cleared with Sepharose beads (Sigma-Aldrich) for 1 h at 4 °C. Anti-CENP-A antibody (CST, 2047) was added to cleared supernatant (total of 10 µg) and incubated for 3 h at 4 °C. The specificity of the antibody was confirmed via RT-PCR against murine centromeric repeats. During primary antibody incubation, beads were blocked in NET Buffer (150 mM NaCl, 50 mM Tris, pH 7.5, 1 mM EDTA, 0.1% IGEPAL and 0.25% gelatin). 100 µl of blocked beads were added per sample and CENP-A ChIP was incubated overnight rotating at 4 °C. Following this, samples were briefly spun and beads were washed three times with Wash Buffer 1 (150 mM NaCl, 20 mM Tris-HCl, pH 8, 2 mM EDTA, 0.1% SDS, 1% Triton X-100), once with High Salt Wash Buffer (500 mM NaCl, 20 mM Tris-HCl, pH 8, 2 mM EDTA, 0.1% SDS, 1% Triton X-100), and the chromatin was eluted 2× each with 200 µl Elution Buffer (50 mM NaHCO3, 0.32 mM sucrose, 50 mM Tris, pH 8, 1 mM EDTA, 1% SDS) at 65 °C for 10 min at 1500 rpm. The elution step was repeated one more time and the two eluates were combined. The input sample was adjusted to a final volume of 400 µl with Elution Buffer. To isolate DNA, 16.8 µl of 5 M NaCl and 1 µl of RNase A (10 mg/ml, Promega) were added and incubated for 1 h at 37 °C. After that, 4 µl of 0.5 M EDTA and 12 µl Proteinase K (2.5 mg/ml, Promega) were added, and samples were incubated for another 2 h at 42 °C. Samples were then subjected to phenol-chloroform extraction followed by purification with a QIAquick PCR Purification column (Qiagen). DNA library was prepared using the NEBNext Ultra II DNA Library Prep Kit for Illumina according to the manufacturer’s instructions. Libraries were sequenced using NovaSeq 6000 paired-end sequencing. A total of 3 replicate experiments were performed. Of the three biological replicates per condition, one replicate per condition (chip_HC_3 and chip_PTZ_3) failed quality control on multiple independent criteria (low inter-replicate correlation, low sequencing depth, and near-absent peak enrichment; Figure 3-figure supplement 1B, Figure 3-figure supplement 1C) and was excluded from all downstream analyses. All subsequent analyses therefore used the two high-quality replicates per condition, with the second condition serving as independent cross-validation.

### Next-generation sequencing data processing

ChIP-seq reads were processed with snakePipes^67^. In short, paired-end raw sequencing reads were trimmed with trim_galore v0.6.5 and aligned to the mouse reference genome GRCm38 using bowtie2 v2.3.5.1. To quantify chromatin binding, alignments were deduplicated using Sambamba v0.7.1 and sequencing coverages calculated via deepTools v3.3.2 over 100-nucleotide windows. To quantify CENP-A binding to centromeres, the trimmed ChIP-seq reads were aligned to a custom reference comprising a head-to-tail trimerized minor satellite consensus (GenBank: X14464.1); a head-to-tail dimerized mouse major satellite consensus (GenBank: V00846.1); and their reverse complements, using Bowtie2 v2.3.5.1, as suggested previously^43^. The global binding patterns of CENP-A to genomic compartments (promoters, introns, etc.) were assessed using ChIPpeakAnno^68^, detecting an overrepresentation in promoters. To explore local binding patterns to specific promoters and gene bodies, CENP-A ChIP coverages were summarized using genomation v1.26 and the mouse transcriptome annotation from UCSC. We quantified CENP-A binding to genomic intervals over pre-defined regions, including gene bodies (defined as regions between transcription start sites and ends, extended 10% up- and downstream) or promoters (defined as the −1 kb, +1 kb regions from the transcription start sites). We then clustered the CENP-A signal over these features using k-means (number of clusters selected by visual inspection; 4 clusters for promoters and 3 for gene bodies), and visualized CENP-A binding patterns as either meta-region line plots, which show the normalized average CENP-A occupancy, or heatmaps, which depict the normalized CENP-A occupancy per individual feature. Because these data are depth-normalized and lack an exogenous spike-in, they report the genomic localization (occupancy) of CENP-A rather than its absolute abundance; absolute CENP-A levels were assessed separately by immunofluorescence and western blot. Aggregated ChIP-seq data are shown from the retained high-quality replicates. To further explore the epigenomic and transcriptional patterns of CENP-A-bound genomic regions, we added public data, including structural features (CpG islands) as well as ChIP-seq genomic coverages to the analysis, while maintaining the genes or promoter clusters driven by CENP-A binding signal alone. We reused publicly available data, either reprocessing them from raw reads using ARMOR (RNA-seq; read alignment with salmon v1.4.0 and differential expression analysis with edgeR v3.36.0) or snakePipes (ChIP-seq; as described above) with default parameters, or reusing their processed data, lifting over to GRCm38 when needed. Data sources: CpGi cpgIslandExt track (UCSC), GSE93011 (GSM2442440, GSM2442457), GSE125068 (GSE125068_ATACseq.tar.gz, GSE125068_ChIPseq.tar.gz, GSE125068_nuRNAseq.tar.gz, SRR8441329, SRR8441330, SRR8441331, SRR8441332, SRR8441333, SRR8441334, SRR8441336, SRR8441337, SRR8441338), GSE85873 (GSM2286402, GSM2286406) and PRJNA390822 (SRR5723788). Code and data are available at https://github.com/imallona/astankovic_cenp_chip and GEO: GSE230784.

### Viral transduction and in vitro neuronal activation

Mature forebrain organoid infection with AAVs and lentiviral vectors: organoids were co-infected with 5 µl of purified viral particles in 50 µl of maturation media. Subsequently, organoids were incubated for 2 h at 37 °C with mild rotation and then returned to standard conditions. Two rounds of infection separated by one week of incubation were performed before activation and fixation. Blue light was delivered using the CoolLED pE-300 excitation system and LNscope (Luigs & Neumann). For light stimulation, single 2.5 ms long, 470 nm pulses at 100% intensity were applied at a frequency of 0.5 Hz for 5 minutes. The frequency and duration of the light pulse were set using a MultiClamp 700B amplifier (Molecular Devices) and pClamp10 (Molecular Devices). The organoids were incubated post-stimulation for 20 minutes in aCSF, fixed and prepared for immunofluorescence. Mouse neural stem cells (80% confluency in 6-well plates) were infected with 4 µl of purified CENP-A-EGFP lentiviral vector and allowed to proliferate for 2 weeks to ensure genomic integration of the transgene. Following this incubation, cells were plated on Laminin-coated (Sigma) coverslips, allowed to adhere overnight and subjected to growth factor removal. Derived neurons were allowed to mature for 2 weeks, followed by depolarization with 90 mM KCl, fixation and immunostaining^62^.

### Fluorescence live cell imaging

All images were acquired using a confocal microscope (LSM 800, Zeiss) fitted with an incubator box, with the temperature set to 37 °C and 5% CO2. Prior to imaging, mNSC-derived neurons were treated with 1 mM TTX for 4 h. Cells were imaged every 3 minutes for 1 h, after which TTX was removed and 90 mM KCl was added, upon which cells were imaged for an additional hour.

### Stereotaxic injections

AAV vectors encoding either control shRNA or shRNA against CENP-A were administered via bilateral stereotaxic injection of the DG^62^. Wild-type C57BL/6 mice were anesthetized by isoflurane inhalation (1% isoflurane for induction and 2-3% isoflurane for maintenance during surgery) and placed in a stereotaxic apparatus (David Kopf Instruments, Tujunga, CA). Prior to surgery, the analgesic Buprenorphine (Buprenorphine hydrochloride, NDC# 12496-0757-5, Reckitt Benckiser, Richmond, VA) was diluted in saline (0.03 mg/ml in sterile 0.9% sodium chloride solution, Vedco Inc., St. Joseph, MO) and injected (0.2 mg/kg, s.c.). For surgery, hair over the skull was removed and a midline incision was made to expose the skull. Injections of virus were made bilaterally (1 µl per hemisphere) using a Hamilton syringe (Cat# 86259, Hamilton, Reno, NV) and a 33-gauge needle. Coordinates were anterior-posterior (AP, relative to Bregma) −2.5 mm, mediolateral (ML, from the midline) ±1.5 mm, and dorsoventral (DV, relative to the skull surface) −2.3 mm.

### Animal behavioral experiments

Behavioral experiments were approved by the Landesdirektion Sachsen in accordance with European and national regulations. Wild-type C57BL/6Rj mice were injected bilaterally with AAVs encoding either control shRNA or shRNA against CENP-A.

Morris Water Maze (MWM)

Mice were tested in the reference memory version of the MWM task as described^46^. Briefly, mice were trained to find a hidden platform in a plastic pool (1.9 m diameter) filled with opaque water. On five consecutive days, six trials per day were performed with an inter-trial interval of at least 60 minutes. In every trial, mice were allowed to search for the platform for 120 s and, irrespective of trial outcome, were guided to the platform position where they stayed for 15 s. Starting positions were changed each day but remained constant throughout individual days for all mice. From day 4 onwards, the platform position was changed to the opposite quadrant. In addition, the first trial of day 4 was a probe trial lasting 60 s, performed without a platform. Analyses of swim paths and search metrics were done based on raw time-tagged xy-coordinates using the RStudio Rtrack package (version 1.0.7) ^46^. In brief, swim paths were classified into non-goal-oriented strategies (thigmotaxis, circling, random path), procedural strategies (scanning and chaining), and allocentric search strategies (directed search, corrected search, direct path and perseverance). For the odds ratio, data were grouped into acquisition (days 1 to 3) and reversal (days 4 to 5) trials, whereas search strategies were grouped into non-spatial, spatial and perseverance. The ratio was calculated based on the odds of CENP-A-knockdown mice using a particular strategy, divided by the odds of control mice using that same strategy. Finally, using the initial path (i.e., initial section of the swim path with the same length as the direct start-to-goal distance) as a reference, initial heading error and initial trajectory error were measured based on the cumulative angles to the direct path and the distance to the goal, respectively. Five mice (3 control and 2 KD) were excluded from the analysis due to their consistent use of the anxiety-related thigmotaxis strategy in more than 33% of trials (compared to less than 20% for the remaining mice).

Novel object recognition task (NORT)

NORT was conducted in an open field arena (50 cm × 50 cm), with three 5-min sessions performed on the same day. In the habituation session, mice were allowed to explore and habituate to an empty arena. After a time interval of 90 minutes, mice were placed back in the arena and presented with two identical objects during the familiarization session. Finally, mice underwent the test session 3 hours later, where they encountered one familiar object and one novel object. Cumulative time of object exploration was measured for familiarization and test sessions, and the discrimination index was calculated as follows: (time with novel object − time with familiar object) / (time with novel object + time with familiar object).

The discrimination index was computed over the first 30 s of object exploration; all 31 mice were included in the analysis.

### Microscopy and image analysis

Confocal imaging of all samples was done using a Zeiss LSM800, controlled with Zeiss ZEN software. All image analyses were performed using FIJI (version 2.9.2) based on ImageJ. Centromeric CENP-A signal was measured as outlined before^69^. In brief, a box was placed around the centroid position of the centromere. The maximum intensity value within the box was corrected for local background by subtraction of the minimum pixel value. As centromeres are highly clustered in post-mitotic neurons, the minimal circularity was set to 0.8 AU. The total nuclear CENP-A fluorescence intensities were calculated by the average Z-projection of imaged slices, followed by measurement of DAPI-masked nuclear CENP-A. To derive fluorescence intensities of non-centromeric CENP-A, an average value of measured centromeres was subtracted from the value measured for the total nuclear CENP-A, per individual cell. The total number of recognized centromeres is comparable across different positions.

### Fluorescence activated cell sorting (FACS)

Nuclei dissociation was performed similarly to a previously described protocol^32^. Briefly, the dentate gyri were microdissected and immediately placed into a nuclei isolation medium (0.25 M sucrose, 25 mM KCl, 5 mM MgCl2, 10 mM Tris-HCl, 100 mM dithiothreitol, 0.1% Triton, protease inhibitors). Tissue was Dounce-homogenized, allowing for mechanical separation of nuclei from cells. Samples were washed, resuspended in nuclei storage buffer (0.167 M sucrose, 5 mM MgCl2, 10 mM Tris-HCl, 100 mM dithiothreitol, protease inhibitors) and filtered. Isolated nuclei were stained for Fos (Santa Cruz, sc-25) and Prox-1 (4G10, EMD Millipore). The nucleic acid stain Hoechst 33342 (5 µM, Life Technologies) was included in the media to facilitate visualization of the nuclei for quantification. Samples were analyzed using an LSR II Fortessa (BD Biosciences) and data were processed using FlowJo (Tree Star).

### Pentylenetetrazol (PTZ) injections

Mice were injected with 45 mg/kg of fresh PTZ dissolved in saline (intraperitoneal; at a volume of 20 ml/kg body weight). Control mice were injected with 400 µl of saline. Mice were sacrificed 1 h after injection. The mice exposed to PTZ exhibited seizure-like symptoms beginning 5 min after PTZ injection^35^. Saline-injected mice did not show any seizure symptoms.

### Quantification and statistical analyses

Statistical analyses were performed in Python (SciPy, scipy.stats, for t-tests and Mann-Whitney tests; statsmodels for linear mixed-effects models, repeated-measures ANOVA, and Holm correction for multiple comparisons), except where noted. The reanalysis of published RNA-seq data in Figure 4-figure supplement 1C was performed in GraphPad Prism 9 (GraphPad Software) using unpaired two-tailed Welch’s t-tests, and the behavioral analyses were performed in R (RStudio; Rtrack version 1.0.7, see Animal behavioral experiments). Unless otherwise stated, the unit of analysis was the biological replicate: the individual mouse for in vivo experiments, the independent differentiation for mNSC-derived neuron cultures, and the cortical unit (with differentiation batch accounted for; see below) for forebrain-organoid experiments. Per-cell or per-centromere measurements were averaged within each biological replicate, and statistical tests were performed on the per-replicate values rather than on pooled individual cells. All t-tests were two-tailed. Exact P values are reported throughout, including for non-significant comparisons; significance is denoted as *, P<0.05; **, P<0.01; ***, P<0.001; ns, not significant. Data are presented as mean ± SD unless stated otherwise, with individual biological-replicate values shown as points.

Fixed-sample image quantification was performed blind to experimental condition. Live-imaging experiments (Figure 2I and Figure 2-figure supplement 1D) were not blinded, as these are continuous single-session recordings in which baseline imaging precedes addition of the depolarizing stimulus (KCl) to the same field of cells; the timing of stimulation is therefore inherent to the acquisition and cannot be masked.

In vivo knockdown experiments comparing shCTRL- and shCENP-A-injected mice followed a randomized block design: each batch contained one shCTRL and one shCENP-A mouse that were injected, and after 4 weeks exposed to the novel environment, and perfused together. These experiments were therefore analyzed as matched pairs using paired two-tailed t-tests (Figures 2F, 2G, 4B and Figure 4-figure supplement 1B), which removes shared batch-to-batch technical variance. Comparisons of FOS-positive versus FOS-negative neurons within the same animals were likewise paired (Figures 1F, 2C, 2D, Figure 1-figure supplement 1H). The FACS quantification of FOS-positive nuclei (Figure 4C), which involved unequal, independent groups (home cage, novel environment, shCTRL, shCENP-A), used unpaired two-tailed t-tests. For the western blot quantification (Figure 1B) and the saline versus PTZ comparison (Figure 1-figure supplement 1F), in which the two groups had markedly unequal variances, an unpaired two-tailed Welch’s t-test was used.

For the time-course experiments in the dentate gyrus (Figure 1E, FOS and non-centromeric CENP-A across home cage, 1 h and 5 h; Figure 1-figure supplement 1K, ARC), independent animals were used at each timepoint (terminal perfusion), and the pre-specified contrasts were compared using unpaired two-tailed t-tests; FOS and non-centromeric CENP-A were quantified independently and were not statistically compared to one another. For the live-imaging experiments in mNSC-derived neurons (Figure 2I, activated; Figure 2-figure supplement 1D, non-activated), each cell’s fluorescence trajectory was normalized to its own first frame and aggregated to a per-replicate session mean (n = 3 independent differentiations per condition). Because first-frame normalization sets the baseline to a constant (removing its variance), the activation response was assessed primarily by comparing the activated and non-activated conditions (unpaired two-tailed Welch’s t-test on per-replicate session means), with a one-sample t-test of the activated per-replicate session means against the normalized baseline of 1.0 as a supporting test. For the KCl time-course in mNSC-derived neurons (Figure 4-figure supplement 1E), normalized CENP-A-EGFP and FOS intensities (non-activated set to 1.0) were analyzed by repeated-measures ANOVA for the overall effect of time on each protein; the difference in kinetics between the two proteins was assessed by a two-way (protein × time) repeated-measures ANOVA. For the chaperone-knockdown screen in mNSC-derived neurons (Figures 2K, 2L), mCherry-positive (shRNA-expressing) and adjacent mCherry-negative cells were compared within each of three biological replicates by paired two-tailed t-tests on per-replicate means, and P values were corrected for multiple comparisons across the five shRNA constructs using the Holm method. The knockdown-validation experiment (Figure 2-figure supplement 1B) and the shCTRL-versus-shDAXX percentage comparisons (Figure 2-figure supplement 1G, H) used paired two-tailed t-tests on per-replicate values.

Forebrain-organoid experiments (Figure 5) used the cortical unit as the biological replicate, with the differentiation batch modelled as a random effect in a linear mixed-effects model, because each batch yielded variable and unequal numbers of usable cortical units per condition (no one-to-one matching across conditions) owing to the stochastic attrition of mature organoids during processing. Between-condition comparisons of shCTRL versus shCENP-A (Figure 5I) were analyzed with batch as a random effect. Where the mixed model was singular or failed to converge, or too few units per batch precluded fitting it (Figure 5B, CTRL versus blue-light-activated; Figure 5H), an unpaired two-tailed Welch’s t-test is reported together with the per-batch concordance of the effect. Comparisons of FOS-positive versus FOS-negative neurons within the same cortical units (Figures 5E, 5F) were paired at the cortical-unit level. In Figure 5, “CTRL” denotes AAV-infected cells not exposed to blue light and “Activated” denotes AAV-infected cells exposed to blue light. Descriptive quantifications with a uniformly near-zero group or a single-group composition (Figures 1D, 5C and Figure 1-figure supplement 1G, J) are reported as observed proportions with the number of cells examined, without an inferential test.

For all imaging quantifications, n values (number of mice, differentiations, or cortical units, and the approximate number of cells examined per replicate) are given in the corresponding figure legends.

For the behavioral experiments, NORT discrimination indices were compared between familiarization and test within each group by two-sample two-tailed t-tests; Morris water maze heading and trajectory errors were compared between groups by unpaired two-tailed t-tests on data binned by day, and strategy usage, latency, path length, velocity and probe trial quadrant preference were analyzed as described in the corresponding figure legends.

### Data and code availability

This paper makes use of existing, publicly available datasets, listed below. CENP-A ChIP-seq data generated in this study have been deposited at GEO under accession GSE230784. All original code is available at https://github.com/imallona/astankovic_cenp_chip. Publicly available datasets were reused either by reprocessing them from raw reads (RNA-seq with ARMOR: read alignment with salmon v1.4.0 and differential expression analysis with edgeR v3.36.0; ChIP-seq with snakePipes) or by reusing their processed data, lifting over to GRCm38 when needed. Data sources: CpGi cpgIslandExt track (UCSC), GSE93011 (GSM2442440, GSM2442457), GSE125068 (GSE125068_ATACseq.tar.gz, GSE125068_ChIPseq.tar.gz, GSE125068_nuRNAseq.tar.gz, SRR8441329, SRR8441330, SRR8441331, SRR8441332, SRR8441333, SRR8441334, SRR8441336, SRR8441337, SRR8441338), GSE85873 (GSM2286402, GSM2286406) and PRJNA390822 (SRR5723788). Any additional information required to reanalyze the data reported in this paper is available from the corresponding authors upon request. Raw tracking data for the Morris water maze experiments (Figure 4D-H and Figure 4-figure supplement 1F-J) are deposited at Zenodo (DOI: 10.5281/zenodo.23327295) and were analyzed with Rtrack version 1.0.7.

## Author Contributions

A.S. performed experiments, analyzed data, conceived the project, and co-wrote the manuscript. D.G.B. performed experiments, analyzed data. I.M., M.D.R. performed and supervised computational analyses. M.G.Q., B.N.J., V.I.K. performed computational analyses of behavioral and scRNA-seq. S.G., A.E.R., A.G., G.K. performed, supervised, analyzed behavioral experiments. N.A.C., C.F. performed, supervised optogentic experiments. S.F. contributed to data interpretation. S.J. conceived the project and co-wrote the manuscript. All authors revised the manuscript.

## Acknowledgements

This work was supported by the Swiss National Science Foundation (TMAG-3_209272, BSCGI0_157859 and 310030_196869 to S.J.), an EMBO Long-term fellowship (to A.S.), an UZH Candoc Fellowship (to D.G.B.), a Boehringer Ingelheim Fond fellowship (to M.G.Q.), the URPP Adaptive Brain Circuits in Development and Learning (AdaBD) of the University of Zurich, and the Zurich Neuroscience Center. We thank Geneviève Almouzni for comments on the manuscript and conceptual input.

---

**Figure 1-figure supplement 1.**
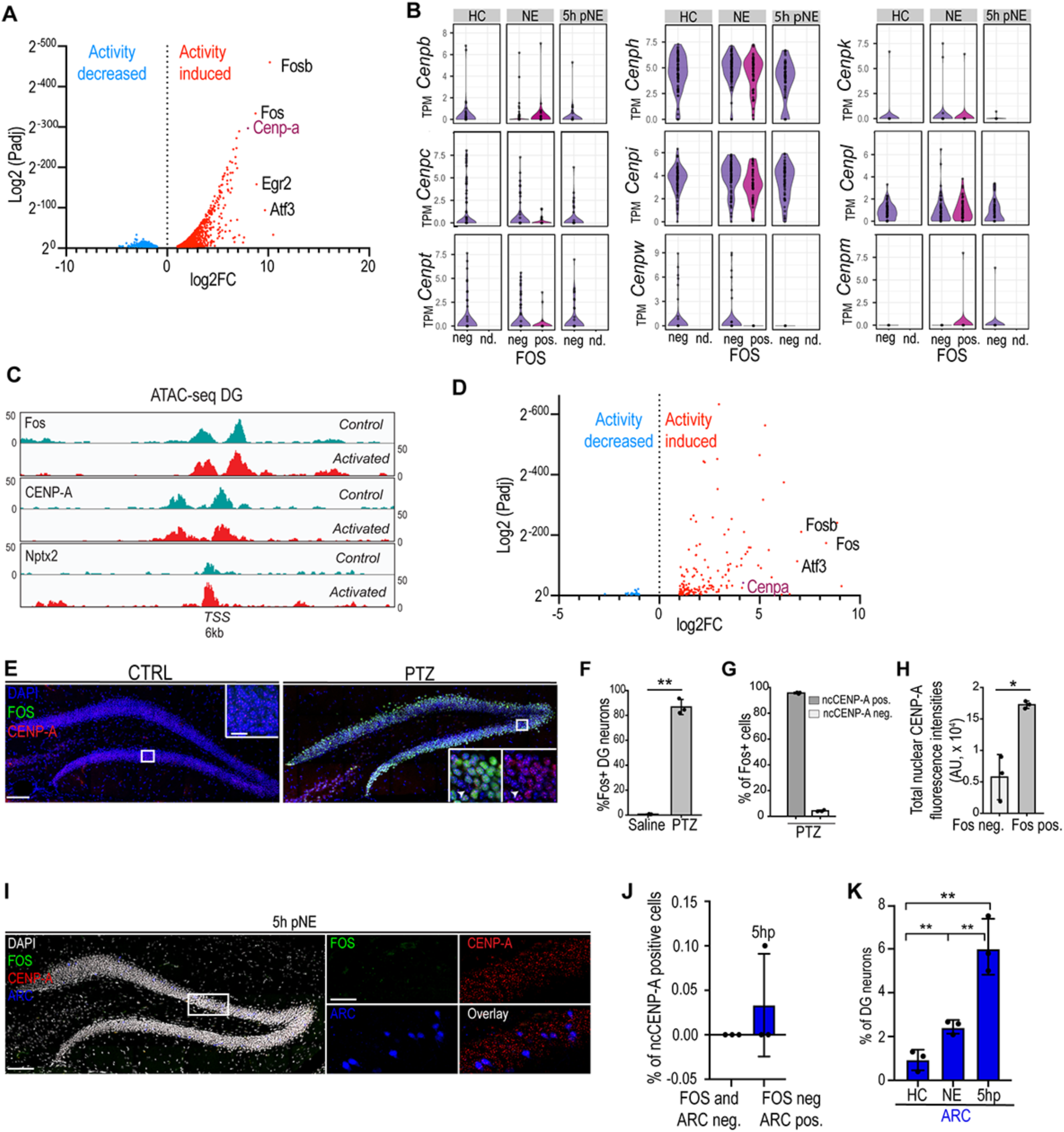
A, Volcano plot of nuclear scRNA-seq coming from a previously published study (17) depicting all significantly down-regulated (activity decreased, blue) or upregulated (activity induced, red) genes present in neurons undergoing activation. Cenp-a mRNA falls within top 5% of overexpressed genes (dark purple). B, Violin plots based on^32^ showing the expression levels of various members of CCAN network in steady and activated neuronal state. C, IGV tracks of ATAC-seq coming from the previously published study^33^. Shown is 6kb window centered around TSS (transcription start site) for control (non-activated, green) and activated (red) granule cells for various IEGs, including CENP-A. D, Volcano plot depicting differentially translated genes in riboRNA-seq, derived from^17^, showing that CENP-A is part of the activity-induced neuronal translatome. E, Pentylenetetrazol (PTZ) induces expression of FOS (green) and CENP-A (red) compared to home cage controls (Saline) (left panel). Boxed areas are shown in high power views. Nuclei were counterstained with DAPI (blue). Scale bars represent 100 µm. F, Percentage of FOS-positive dentate gyrus neurons in saline-treated versus PTZ-treated mice. PTZ induced widespread FOS (∼0.68% vs ∼86.7%). N=3 mice per group; unpaired two-tailed Welch’s t-test, P=0.0016 (**). Points are per-mouse percentages. G, Percentage of FOS-positive neurons that are also non-centromeric-CENP-A-positive, in PTZ-treated mice (co-localization within the PTZ condition). Mean 95.6% (1026/1073 cells; per mouse 96.1, 96.0, 95.0%). N=3 mice; descriptive. Points are per-mouse percentages. H, Total nuclear CENP-A fluorescence intensities in FOS-positive versus FOS-negative neurons within PTZ-treated mice. Nuclear CENP-A was higher in FOS-positive neurons. N=3 mice; paired two-tailed t-test, P=0.025 (*). Points are per-mouse means. I, Mice were exposed to Novel environment (NE) exploration for 1h and then returned to Homecage for 5h. Sections were stained with IEG FOS (green), CENP-A (red) and ARC (blue). Nuclei were counterstained with DAPI (white). Scale bar represents 50 µm. J, Non-centromeric CENP-A at 5 h post novel-environment, in double-negative and ARC-positive neurons. Non-centromeric CENP-A was essentially absent at this timepoint (0 of ∼50 cells per condition per mouse in most animals). N=3 mice; descriptive. Points are per-mouse percentages. K, Percentage of ARC-positive dentate gyrus neurons at home cage (HC), 1 h and 5 h post novel-environment. ARC rose and remained elevated (HC 0.93%, 1 h 2.45%, 5 h 6.10%; HC vs 1 h P=0.010; HC vs 5 h P=0.003; 1 h vs 5 h P=0.009), a sustained/late profile that contrasts with the transient FOS and non-centromeric CENP-A dynamics (Figure 1E). N=3 mice per timepoint; unpaired two-tailed t-tests. Points are per-mouse percentages. (*, P<0.05, ** P<0.01, *** P<0.001). Figure 1-figure **supplement 1-source data 1.** FOS induction after PTZ. Figure 1-figure **supplement 1-source data 2.** Non-centromeric CENP-A in FOS-positive neurons after PTZ. Figure 1-figure **supplement 1-source data 3.** Total nuclear CENP-A after PTZ. Figure 1**-figure supplement 1-source data 4.** Non-centromeric CENP-A at 5 h after novel environment. Figure 1-figure **supplement 1-source data 5.** ARC time course.

**Figure 2-figure supplement 1.**
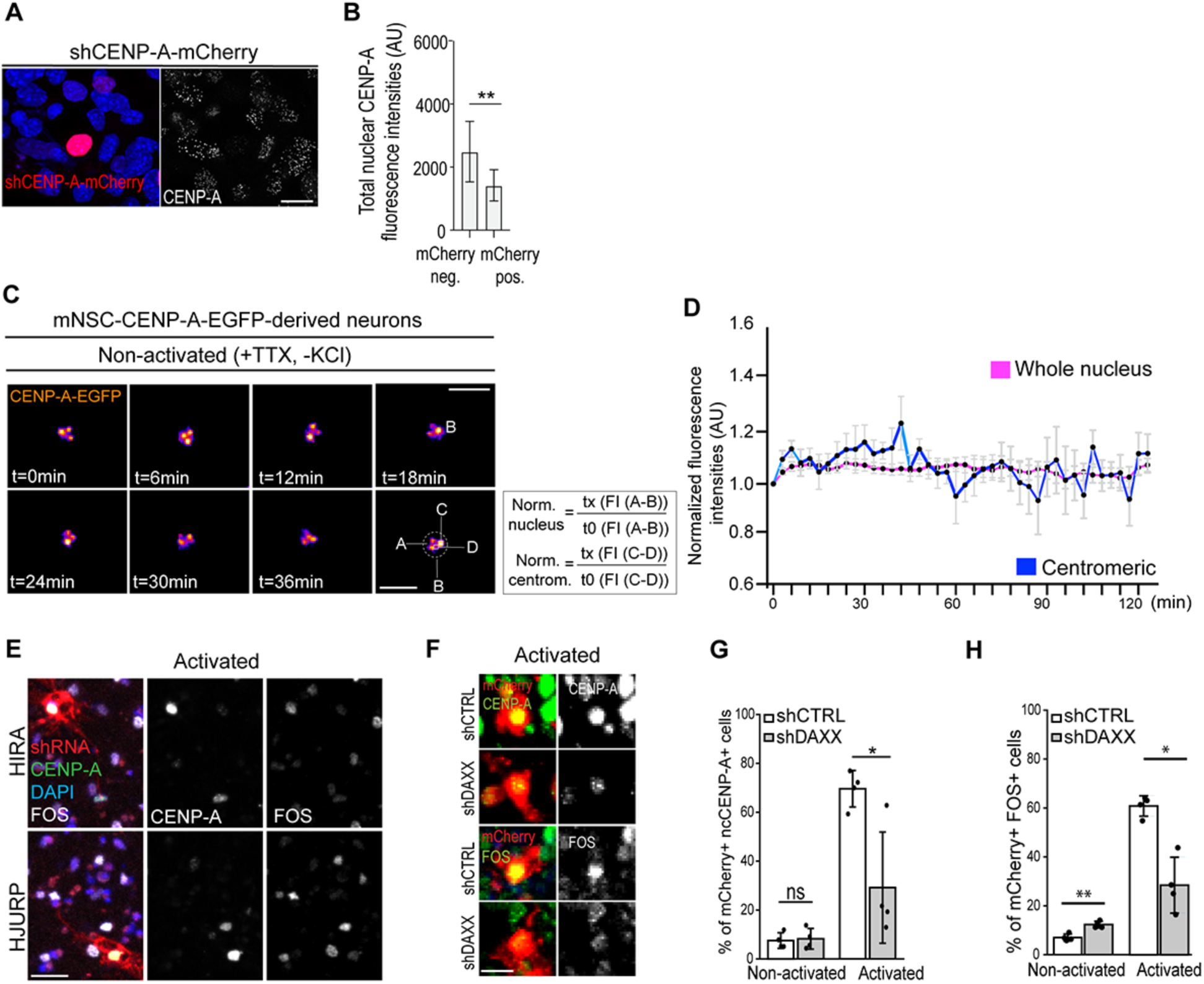
A, CENP-A (white) levels were assayed 72h post-electroporation with construct harboring shCENP-A-mCherry (red) in proliferating mouse neural stem cells (mNSCs). Scale bar denotes 5 µm. B, Total nuclear CENP-A fluorescence (background-corrected) in mCherry-positive (hairpin-expressing) versus mCherry-negative cells, following electroporation of proliferating mNSCs with the CENP-A hairpin used in vivo. The hairpin reduced CENP-A by 44% (1.79-fold). N=3 replicates (∼10 cells per condition per replicate); paired two-tailed t-test on per-replicate means, P=0.009 (**). Points are per-replicate means. C, Shown are TTX (tetrodotoxin)-treated stills of live-imaged CENP-A-EGFP mNSC-derived neurons. Sample was imaged every 3 mins, shown are each second frame. D, Live imaging of CENP-A-EGFP in non-activated mNSC-derived neurons over 42 frames (∼3 min per frame). Lines and markers, mean of 3 independent differentiations; bars, SD across replicates; whole nucleus versus centromeric. Neither measure shows the rise seen upon activation (Figure 2I). Between-condition comparison (activated, Figure 2I, versus non-activated; unpaired two-tailed Welch’s t-test): whole nucleus P=0.02; centromeric P=0.20 (ns). N=3 biological replicates. E, Representative images of mNSC-CENP-A-EGFP-derived neurons treated with the indicated shRNA constructs undergoing KCl-induced activation. Scale bar denotes 5 µm. F, Representative images of mNSC wild-type-derived neurons treated with the indicated shRNA constructs with (A; activated) or without (NA; Non-activated) KCl stimulation. Cells were counterstained for CENP-A and FOS. Scale bar denotes 5 µm. G, Percentage of mCherry-positive (shRNA-expressing) cells that are non-centromeric-CENP-A-positive, for shCTRL and shDAXX, in non-activated (NA) and activated conditions. Without activation the fraction did not differ (7.5% vs 8.2%; P=0.81, ns). Upon activation, DAXX knockdown reduced the fraction (69.6% vs 29.2%; P=0.046; lower in all four replicates). N=4 biological replicates; paired two-tailed t-tests. Points are per-replicate percentages. H, Percentage of mCherry-positive (shRNA-expressing) cells that are FOS-positive, for shCTRL and shDAXX, in non-activated (NA) and activated conditions. DAXX knockdown slightly elevated the FOS-positive fraction at baseline (7.1% vs 12.3%; P=0.003) but strongly reduced the activation-induced FOS response (60.8% vs 28.4%; P=0.012; lower in all four replicates). N=4 biological replicates; paired two-tailed t-tests. Points are per-replicate percentages. (*, P<0.05, ** P<0.01, *** P<0.001). Figure 2-figure **supplement 1-source data 1.** Knockdown validation. Figure 2-figure **supplement 1-source data 2.** Live imaging, non-activated, per-cell traces. Figure 2-figure **supplement 1-source data 3.** Non-centromeric CENP-A after DAXX knockdown. Figure 2-figure **supplement 1-source data 4.** FOS after DAXX knockdown.

**Figure 3-figure supplement 1.**
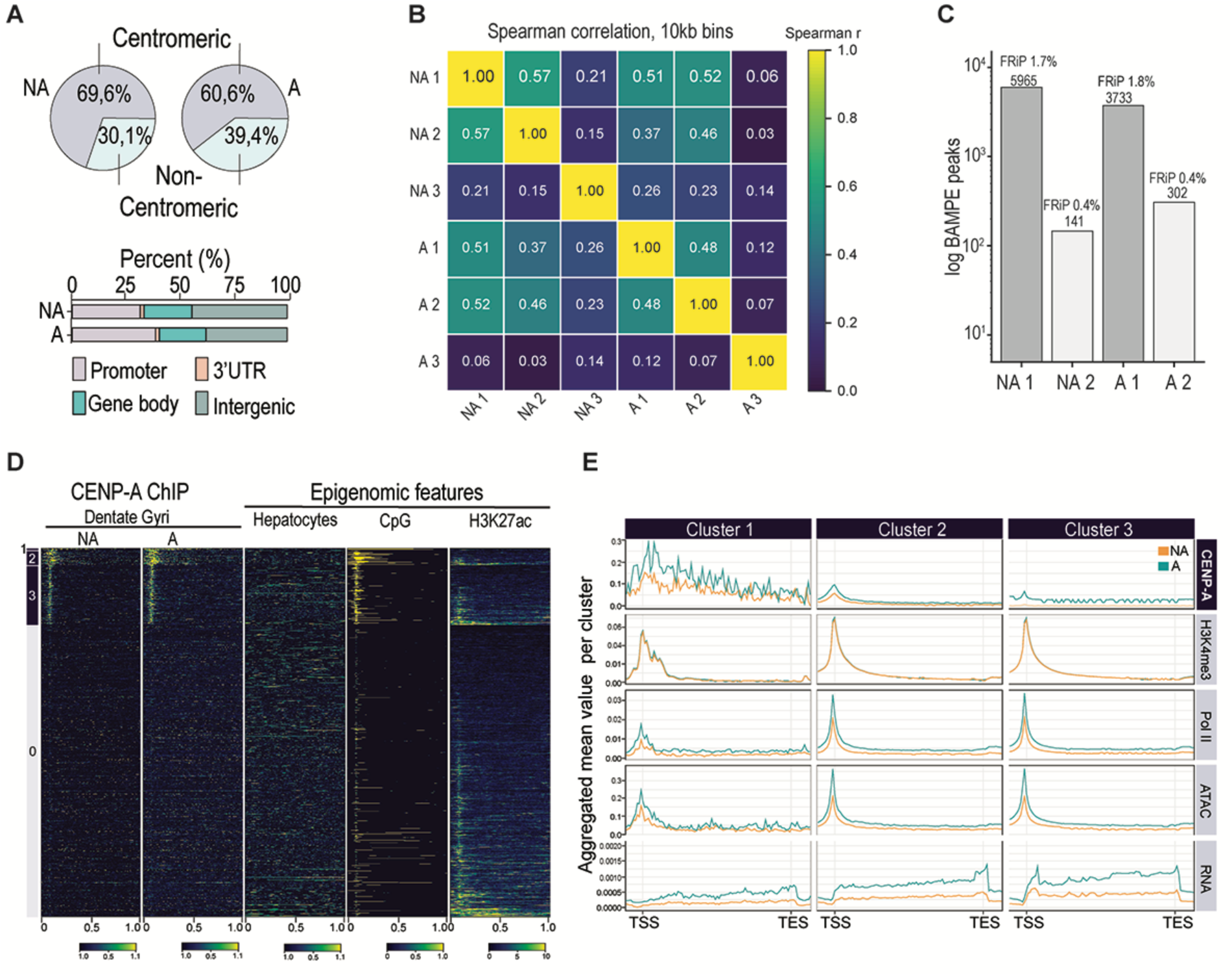
A, Proportion of CENP-A ChIP reads at centromeric versus non-centromeric regions, in non-activated and activated states. Depth-normalized read-composition fractions; the apparent NA/A difference reflects read normalization and is not a quantitative between-state comparison of abundance. B, Spearman correlation of CENP-A ChIP read coverage in 10 kb genome bins across all libraries. Replicate 3 of each condition is a correlation outlier (shown to justify its exclusion). Replicate 3 present. C, Number of MACS2 (BAMPE, broad) peaks per library on assembled chromosomes (log scale), annotated with fraction of reads in peaks (FRiP). Replicate 3 excluded on QC criteria (absent from this panel; see Methods). D, CENP-A occupancy over gene bodies in non-activated (NA) and activated (A) states, grouped into three gene-body clusters (plus a background group), with reused epigenomic features. NA and A shown to demonstrate conserved occupancy. These are gene-body clusters (3), distinct from the promoter clusters (4) in Figure 3 (see Methods). E, Aggregated mean signal per gene-body cluster for CENP-A and reused marks (H3K4me3, Pol II, RNA), non-activated versus activated, across the three gene-body clusters. As in Figure 3B, the NA/A traces for the epigenomic/RNA marks are external datasets (see Methods). Figure 3-figure **supplement 1-source data 1.** ChIP-seq library correlation and PCA. Figure 3-figure **supplement 1-source data 2.** Peak counts and FRiP.

**Figure 4-figure supplement 1.**
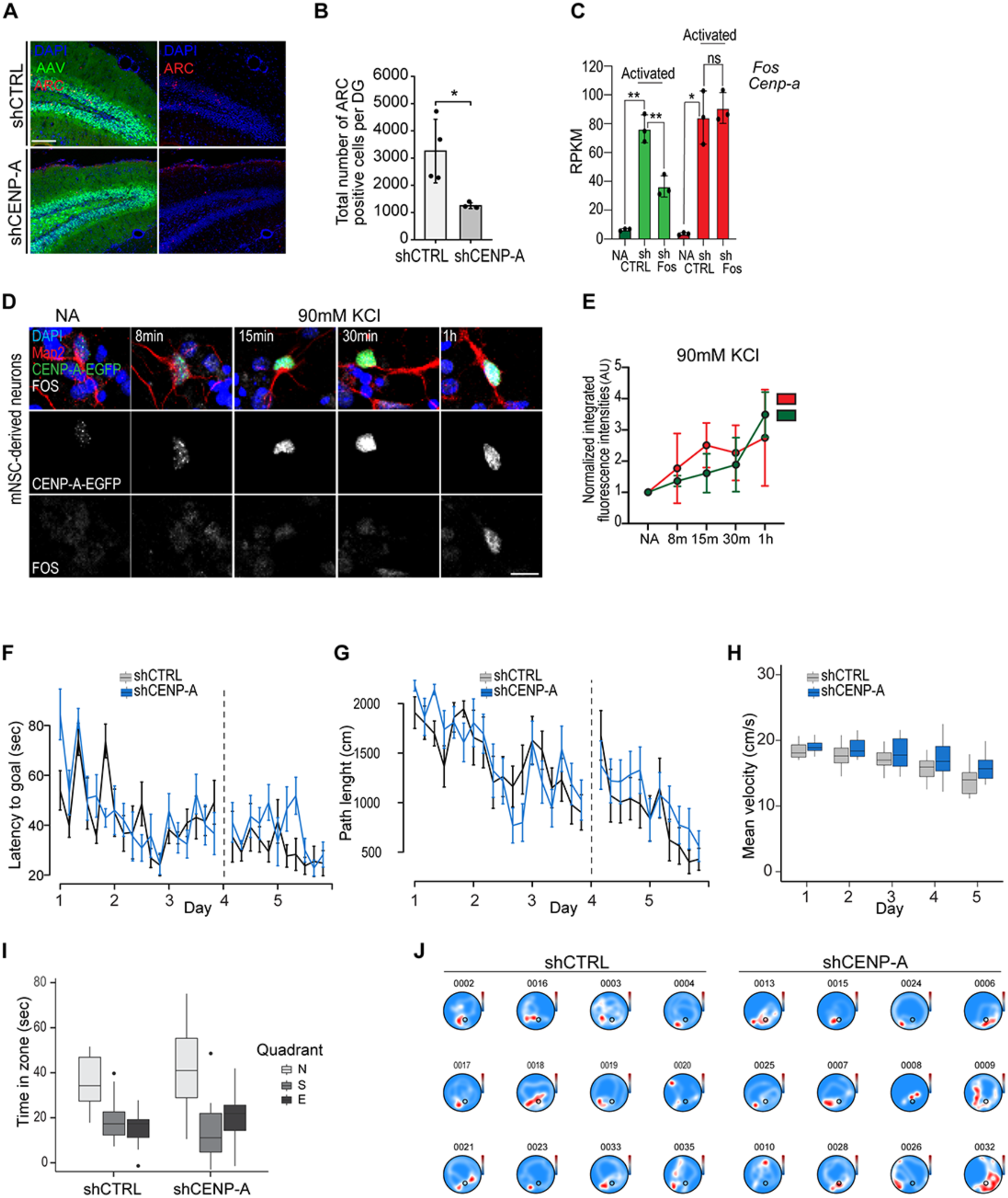
A, AAV-mediated knockdown of CENP-A (shCTRL, left panel; shCENP-A, right panel; green) reduces number of ARC-labeled cells (red) after NE. Nuclei were counterstained with DAPI (blue). Scale bar represents 50 µm. B, Total number of ARC-positive cells per dentate gyrus in shCTRL- and shCENP-A-injected mice. Each batch contained one shCTRL and one shCENP-A mouse processed together; data are analysed as matched pairs. shCENP-A reduced ARC-positive cells ∼2.6-fold (the knockdown mouse lower than its matched control in every batch). N=4 matched pairs (mice); paired two-tailed t-test, P=0.035 (*). Points are per-animal counts. This parallels the reduced FOS response (Figure 4). C, Bulk RNA-seq expression (RPKM) of Fos (green) and Cenpa (red) in non-activated and activated dentate gyrus, in the background of shCTRL or shRNA against Fos; reanalysis of published data (see Methods). Fos knockdown reduced Fos expression 2.1-fold upon activation (P=0.006), confirming effective knockdown, but did not affect Cenpa expression (1.08-fold; P=0.63, ns), indicating activation-induced Cenpa expression is independent of Fos. N=3; unpaired two-tailed Welch’s t-tests. D, *In vitro* activation of mouse neural stem cell (NSCs)-derived neurons. Neurons were counterstained for Map2beta (red) and FOS (green). Nuclei were counterstained with DAPI (blue). Scale bar represents 5 µm. E, Normalized CENP-A-EGFP (red) and FOS (green) fluorescence intensities in the same mNSC-derived neurons at 8, 15, 30 and 60 min after KCl stimulation; non-activated intensities set to 1 for both proteins. Both proteins were significantly induced over the time course (repeated-measures ANOVA: CENP-A P=0.004; FOS P=0.001). CENP-A rose rapidly and appeared to plateau by ∼15 min, whereas FOS continued to accumulate through 1 h; this apparent difference in kinetics did not reach statistical significance (protein x time interaction P=0.23). N=3 biological replicates (∼60 cells per timepoint per replicate); points are per-replicate means, lines are means, error bars SD across replicates. F-H, Group effects during learning and re-learning were assessed using repeated measurements ANOVA (Latency to goal: d1-d3: p = 0.361; d4-d5: p < 0.01; Path length: d1-d3: p = 0.901; d4-d5: p = 0.349, Velocity: d1-d3: p < 0.01; d4-d5: p < 0.01). Analyses for group effects on individual days were performed using Wilcoxon signed-rank test (Latency to goal on d4: p=0.091, Path length on d4: p = 0.248, Velocity on d4 < 0.01). All animals learned the task, indicated by repeated measurements ANOVA showing a significant effect of days of training (Latency to goal: d1-d5: p < 0.0001, Path length: d1-d5: p < 0.0001, Velocity: d1-d5: p < 0.05). I, Quadrant preference on probe trial was evaluated using ANOVA, with Bonferroni p-adjustment. \**p* < 0.05; \*\**p* < 0.01; \*\*\**p* < 0.001; n.s., non-significant. J, Individual heatmaps for probe trial are shown for shCTRL (left) and shCENP-A (right)-injected mice. (*, P<0.05, ** P<0.01, *** P<0.001). Figure 4-figure **supplement 1-source data 1.** ARC-positive cells after knockdown. Figure 4**-figure supplement 1-source data 2.** Fos and Cenpa expression after Fos knockdown. Figure 4-figure **supplement 1-source data 3.** CENP-A-EGFP and FOS time course after KCl. Figure 4-figure **supplement 1-source data 4**. Morris water maze raw tracking data (deposited at Zenodo, DOI: 10.5281/zenodo.23327295); same dataset as Figure 4-source data 3.

**Figure 5-figure supplement 1.**
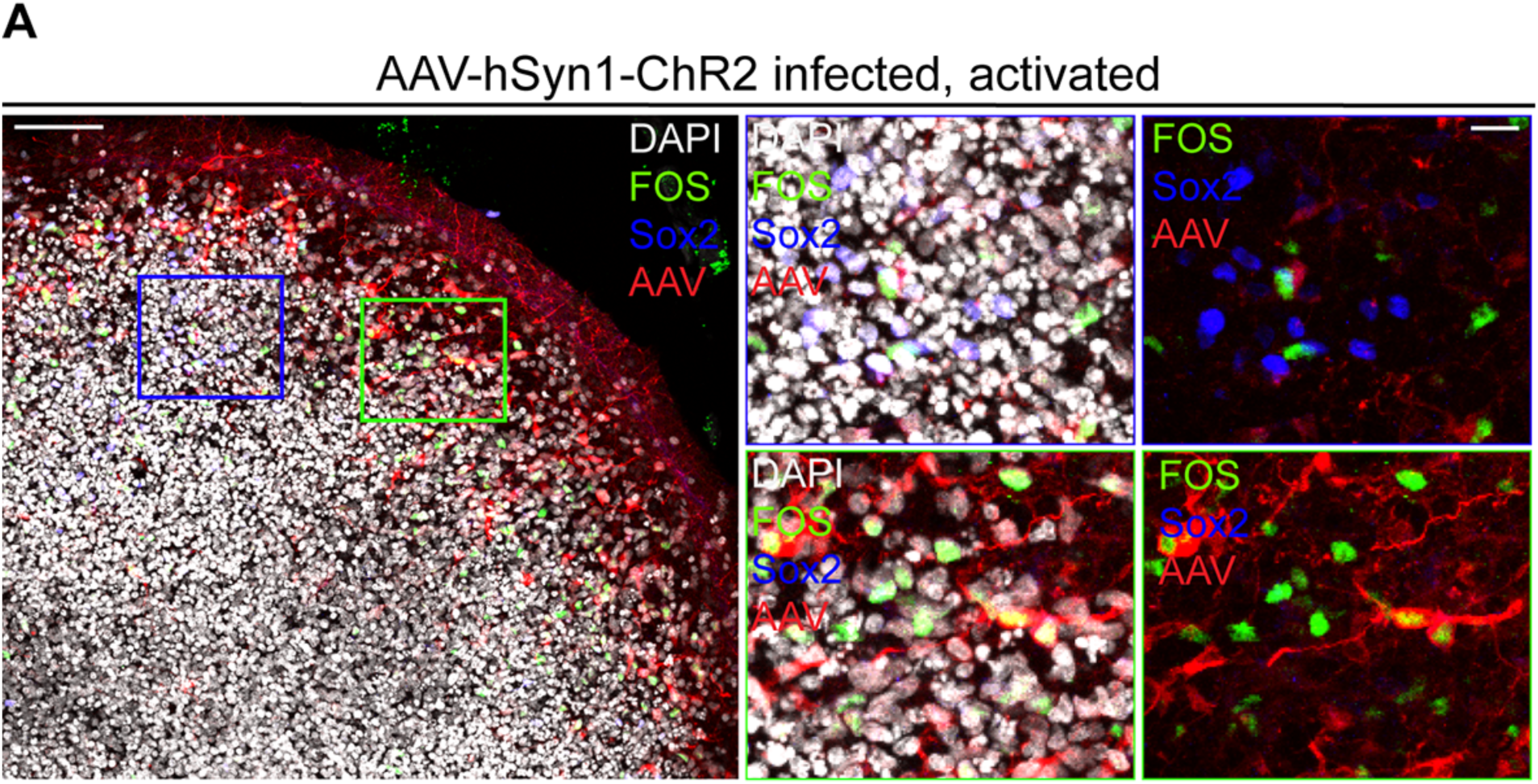
A, Representative examples of an organoid infected with AAV-hSyn1-ChR2 (red) under activated conditions (exposure to blue light). Activated neurons are labeled with FOS (green) and neural progenitor cells with SOX2 (blue). Boxed areas are shown in high power views. Scale bars represent 100 µm and 50 µm, respectively.

